# Consensus Pituitary Atlas, a scalable resource for annotation, novel marker discovery and analyses in pituitary gland research

**DOI:** 10.1101/2025.10.28.685060

**Authors:** Bence Kövér, Thea L. Willis, Olivia Sherwin, James Kaufman-Cook, Yasmine Kemkem, Miriam Vazquez Segoviano, Emily J. Lodge, Michel Zamojski, Natalia Mendelev, Zidong Zhang, Gregory R. Smith, Daniel J. Bernard, Hui-Chun Lu, Stuart C. Sealfon, Frederique Ruf-Zamojski, Cynthia L. Andoniadou

## Abstract

Previous single-cell profiling studies of the pituitary gland have yielded minimally reproducible insights largely due to their low statistical power and methodological inconsistencies. To address this problem, we generated a uniformly pre-processed Consensus Pituitary Atlas (CPA) using all existing mouse pituitary single-cell datasets (267 biological replicates, >1.1 million high-quality cells). The CPA revealed novel cell typing and lineage markers, including low-expression transcripts that previous analyses could not detect. The scale of the CPA enabled the development of machine learning models to automate and standardize cell type annotation and doublet identification for future studies. Leveraging the curated metadata, we identified sex-biased and age-dependent gene expression patterns at cell type resolution. To identify drivers of cell fates, first we determined consensus cell communication patterns. Secondly, we used RNA-sequencing and chromatin accessibility data to identify transcription factors associated with cell fates across modalities. The *epitome* platform acts as an interface with the CPA, allowing streamlined user-friendly analyses.

**Highlights:**

- Uniform processing of 267 mouse pituitary single-cell datasets (>1.1M cells)
- The statistical power enabled cell type, sex- and age-specific marker discovery
- Machine learning models facilitate doublet detection and cell typing in new datasets
- *epitome* platform provides programming-free data access and visualizations

## Introduction

The pituitary is a key endocrine gland that regulates major hormonal axes across vertebrates, comprising anterior and posterior pituitary lobes. The anterior pituitary originates from an oral ectoderm invagination termed Rathke’s pouch,^1^ composed of SOX2+ progenitor cells.^2,3^ These progenitors self-renew and give rise to all endocrine pituitary cell types.^4^ Postnatally, SOX2+ cells persist and retain these properties, establishing them as pituitary stem cells.^4^ During differentiation, the stem cells commit to three lineages, marked by POU1F1 (PIT1),^5^ TBX19 (TPIT)^6^ or NR5A1 (SF1).^7^ POU1F1+ cells give rise to thyroid-stimulating hormone-producing thyrotrophs, prolactin-producing lactotrophs, and growth hormone-producing somatotrophs. The NR5A1+ progenitors differentiate into follicle-stimulating hormone and luteinising hormone-producing gonadotrophs. Finally, the TBX19+ lineage gives rise to adrenocorticotropic hormone-producing corticotrophs and the melanocyte-stimulating hormone-producing melanotrophs of the intermediate lobe. The posterior pituitary, of neural origin, releases the hypothalamic hormones oxytocin and vasopressin/antidiuretic hormone and contains pituicytes. In addition to endocrine and neuroendocrine lineages, the gland includes endothelial, mesenchymal, and immune cells.^8^

To characterize pituitary cellular heterogeneity, single-cell and single-nucleus RNA-sequencing and Assay for Transposase Accessible Chromatin (ATAC) studies have been performed on 265 samples^8–43^ profiling over 1.1 million cells (**Figure 1A**). These studies differ in sample preparation, sequencing technologies, read alignment, and downstream analyses, all of which affect outcomes.^44^ Consequently, pituitary single-cell studies report varied numbers of cell types and sub-clusters,^13,33,34,37^ including non-replicable populations such as multihormonal^12^ or “unknown” cells.^40^ Despite abundant data, consensus on pituitary cell type markers remains limited beyond a few canonical examples.

**Figure 1.**
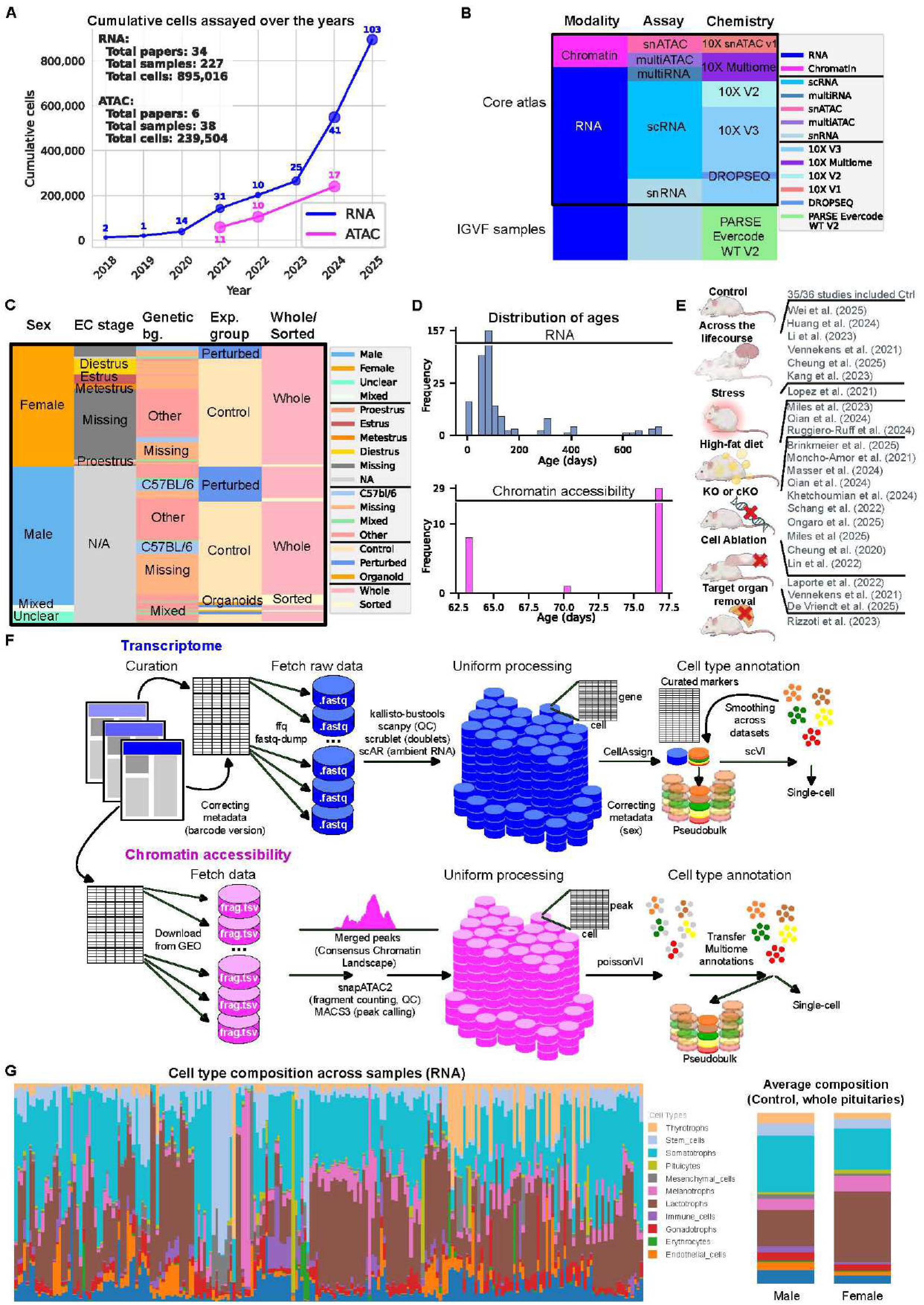
A census of single-cell experiments and generation of the Consensus Pituitary Atlas. (A) Cumulative number of cells profiled (RNA in blue, ATAC in pink) up to Oct 23, 2025. Numbers above dots show sample counts; dot size reflects publication count. (B) Metadata summary showing modality, assay type, and chemistry version per sample. (C) Metadata of biological variables: sex, estrous cycle, genotype, condition (control/perturbed or organoid), and tissue type (whole/sorted). (D) Histogram of sample age distribution (days) for transcriptomic (top) and chromatin accessibility (bottom) samples. Note the broken Y-axes. (E) Schematic of various experimental strategies from the included studies. (F) Schematic of the processing workflow from raw to processed datasets for transcriptomic (top, blue) and chromatin accessibility (bottom, purple) samples. (G) Cell type composition across all RNA samples (left) and mean composition by sex (right).

Uniform data processing improves consistency in bulk^45,46^ and single-cell RNA-seq analyses,^47,48^ and, when applied across pituitary datasets, can resolve these inconsistencies. To this end, we created the Consensus Pituitary Atlas (CPA), encompassing all publicly available samples as of October 2025. The CPA enables high-fidelity cell typing and lineage marker identification, and reveals cell type-specific age and sex effects. We further leveraged the CPA to train bespoke models for doublet detection and cell type annotation, validated on a new postnatal day 4 (P4) male multiome sample. The CPA also identifies transcription factors associated with cell fates as well as ligand-receptor interactions, validated by mRNA *in situ* hybridization and immunofluorescence. To facilitate access and visualization, we developed *epitome* (electronic pituitary omics), a public platform for hypothesis generation, publication-ready figures, and democratized access to uniformly pre-processed pituitary datasets.^49^

## Results

### Generating the Consensus Pituitary Atlas (CPA)

To generate the CPA, all publicly available datasets as of October 28, 2025 were curated, comprising 200 single-cell mouse pituitary samples (**Figures 1B, C**).^8–43^ The curation process revealed and corrected several metadata errors: 13 of 35 publications or their deposited data contained inaccuracies, including incorrect 10X barcoding kit (13/200 samples),^16,17,22,29,31,36^ wrong^17,18,42^ or missing^25,26,31,39,43^ sex information (16/200 samples) and mixed up sample metadata (2/200 samples)^23^ (**Supplementary Table 1**). Gene expression datasets spanned a broad age range, while chromatin accessibility datasets derived exclusively from 7-11 week-old animals (**Figure 1D)**. The datasets included control samples (145/200 samples), genetic, dietary and surgical perturbations (20 perturbations across 51 samples) and 4 stem cell organoid samples (**Figure 1E**).

After uniform pseudoalignment, processing and quality control (**Figure 1F**), three datasets (two generated with early DROPSEQ technology) were excluded from further analyses. To the remaining 197 core samples, we added a recently published set of similarly pseudoaligned datasets from the Impact of Genetic Variation on Function (IGVF) project,^41^ generated with PARSE Biosciences technology (**Figure 1B**), contributing an additional 57 samples after quality control. Cell types were annotated using established marker criteria (see Methods).

Uniform processing of chromatin accessibility data generated a Consensus Chromatin Landscape (CCL), a set of peaks independently identified in at least 10 datasets (**Supplementary Table 2**). Using CCL-derived peaks increased the average fragment count per dataset to 48.5% of all fragments, compared with 43.7% using dataset-specific peaks (**Supplementary Figure 1A**). These CCL peaks capture pituitary cell types similarly, regardless of cell type-abundance bias (**Supplementary Figure 1B**). The CCL can complement peak-calling in future studies, and enable more informative analyses of chromatin accessibility. Cell type annotations were transferred from transcriptomic data using cells assayed in both modalities (multiome assay). Following annotation, all known cell types appeared in their expected proportions across RNA and chromatin modalities (**Figure 1G; Supplementary Figure 1C**).

Considering both modalities, the atlas comprises 1,133,668 high-quality cells (1,020,914 from the core atlas and 112,754 from IGVF samples), derived from 254 biological replicates out of 265 initially assayed.

### Markers enable doublet detection and scalable cell type annotation

To address persistent inconsistencies in pituitary cell type labeling, we reasoned that high-quality annotated data could be used to derive robust cell typing markers and train a cell typing model. To expand the repertoire of cell type markers, we identified highly differentially expressed genes (DEGs) by pseudobulking cells profiles, as pseudobulk statistical approaches outperform single-cell-level methods.^50–56^ Canonical markers (e.g. hormone genes) previously used for annotation were excluded, retaining only novel genes, shown in **Figure 2A** and **Supplementary Table 3**. Identified markers were then used to develop doublet detection and cell type annotation models.

**Figure 2.**
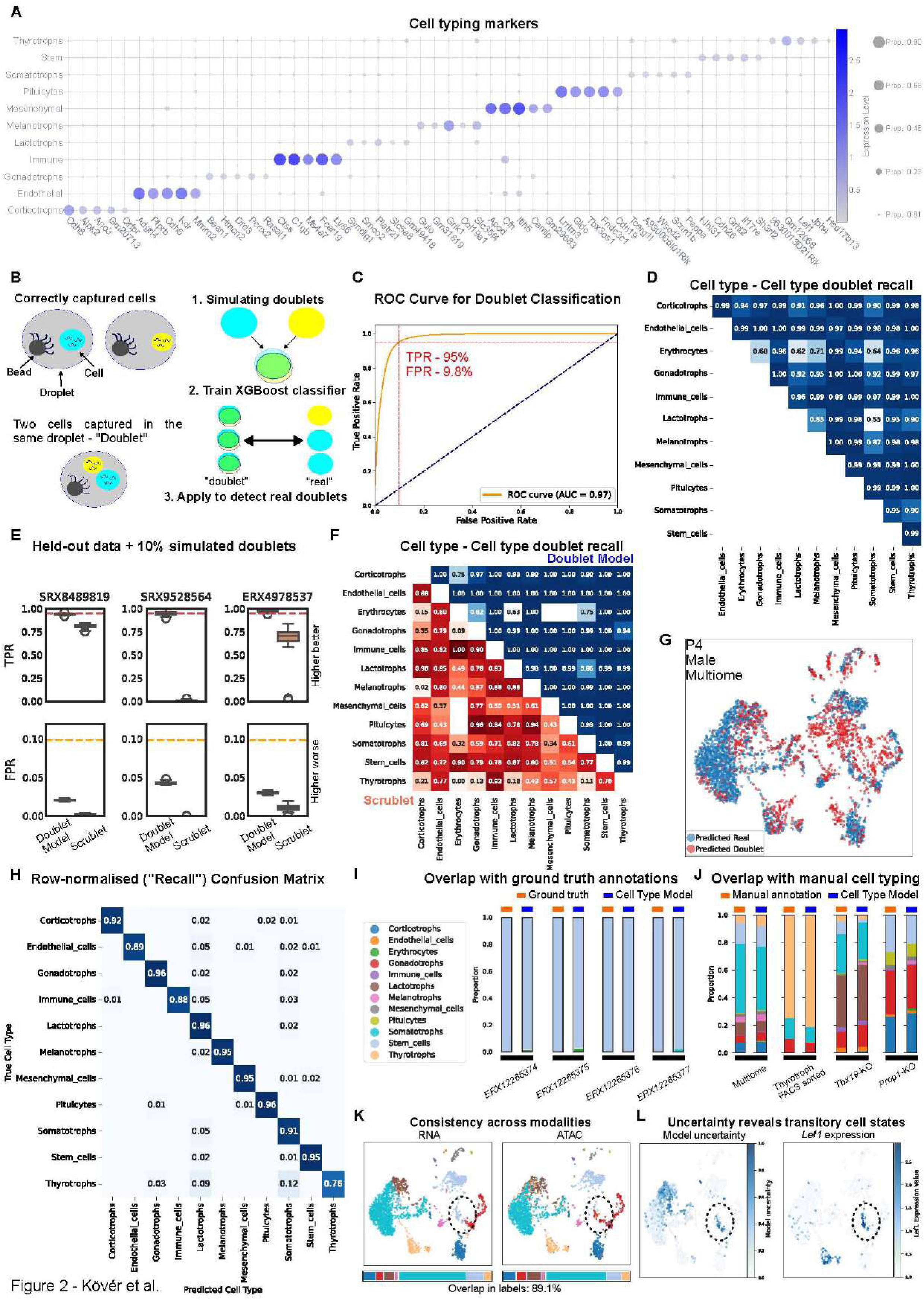
Machine learning models accurately detect doublets and annotate cell types. (A) Dot plot of the top 5 cell typing markers from each cell type. (B) (Left) Schematic showing the composition of droplets in droplet-based single-cell RNA-sequencing, which might contain single cells or two or more cells (doublets). (Right) High-quality filtered and annotated cells were used to simulate doublets (1), and train an XGBoost classifier to separate doublets from real cells (2), before applying to unseen datasets for doublet detection (3). (C) Receiver operating characteristic curve of the Doublet Model from the first fold of a 5-fold cross-validation. (D) Recall matrix for simulated doublets of given pairs of cell types. Color intensity of tiles is proportional to the percentage shown. (E) Box plots comparing true and false positive rates (TPR, FPR) of the Doublet Model and Scrublet across three held-out datasets containing 9.8% simulated doublets. Red and orange dashed lines indicate calibrated thresholds (95% TPR, 9.8% FPR). (F) Recall matrix for simulated doublets in evaluation datasets, with the Doublet Model (blue) above and Scrublet (orange) below the diagonal. Blank tiles denote combinations not observed in simulations. (G) UMAP of the new P4 male multiome dataset showing predicted doublet cells (red) and real cells (blue). (H) Row-normalized confusion matrix comparing Cell Type Model predictions (x-axis) to true labels (y-axis), derived from transcriptomic data. (I) Comparison of predicted and ground-truth labels for stem cell organoid samples.^37^ (J) Comparison of predicted and manually assigned labels for the new P4 male multiome dataset and three held-out samples.^9,13,15^ (K) UMAPs of the P4 male multiome dataset colored by RNA- and ATAC-based cell type predictions (top) and corresponding predicted cell type composition (bottom). (L) UMAPs of the same dataset colored by model uncertainty (left) and *Lef1* expression (right).

Doublets, artefactual pairs of cells captured in the same droplet, complicate cell type annotation (**Figure 2B**). We hypothesized that a pituitary-specific model would outperform generic approaches. Accordingly, we developed the pituitary-specific “Doublet Model” (DM) using the XGBoost gradient boosting framework^57^ (**Figure 2B**; see Methods). Upon 5-fold cross-validation, the DM achieved 86-92% accuracy in distinguishing true cells from doublets across unseen studies. In the first validation fold, the area under the receiver operating characteristic curve (AUROC) was 0.97 (**Figure 2C**). The prediction threshold was calibrated to yield a 95% true positive rate (TPR) at a 9.8% false positive rate (FPR) i.e. the model correctly identified 95% of doublets, while misclassifying only 9.8% of real cells (**Figure 2D**).

To benchmark the DM, we compared its performance with the commonly used Scrublet algorithm^58^ applied during the initial workflow (**Figure 1C**). Three high-quality datasets^9,13,15^ were excluded from training and used for benchmarking. For each, DM and Scrublet were evaluated across 20 simulations containing 10% doublets. The DM consistently outperformed Scrublet (**Figure 2E**), achieving approximately 95% TPR and only ∼2-5% FPR across all datasets. Scrublet repeatedly failed to detect simulated doublets in one dataset (SRX9528564) and failed to identify many doublet combinations in the remaining two (SRX8489819, ERX4978537), whereas the DM successfully detected them (**Figure 2F**). These results highlight the superior accuracy of a tissue-specific model, which can be adapted to other organs.

To demonstrate doublet filtering and subsequent cell typing on new data, we generated a single-nucleus multiome dataset from a postnatal day 4 (P4) male mouse pituitary. DM predicted 33.7% of cells as doublets (**Figure 2G**). These showed higher transcript counts and numbers of detected genes than predicted real cells (**Supplementary Figure 2A**), supporting the validity of the model’s predictions.

We next trained a “Cell Type Model” (CTM) using XGBoost, which achieved 90-95% accuracy across unseen studies in five-fold cross-validation. All cell types reached high recall (88-96%; **Figure 2H**), except thyrotrophs, which showed 76% recall. As a first evaluation, the CTM was applied to 4 pituitary stem cell organoid samples,^37^ showing 97-99.7% concordance between predicted and ground-truth stem cell annotations (**Figure 2I**). As a second evaluation, the P4 multiome data and three other held-out samples were tested: one thyrotroph-enriched,^23^ one *Tbx19* knock-out^34^ and one *Prop1* knock-out.^29^ CTM annotations showed 81-89% agreement with unbiased manual annotations (Methods; **Figure 2J**), confirming accurate detection of enriched or absent cell types in the respective datasets. Notably, the CTM identified a subset of immature melanotrophs (*Pomc* negative) in the *Tbx19*-KO dataset,^34^ which manual annotation had missed (**Supplementary Figure 2B**).

Following this, the CTM was trained on chromatin accessibility data, achieving 92-93% accuracy across unseen studies in three-fold cross-validation (**Supplementary Figure 2C**). All cell types showed high recall (85-97%), except thyrotrophs, which were sometimes misclassified as somatotrophs or lactotrophs (66% recall). Evaluation on the P4 multiome dataset showed 89.2% concordance between chromatin-based and transcriptomic cell type predictions (**Figure 2K**). After annotation, the newly generated P4 sample was added to the CPA, bringing the total to 256 biological replicates and 1,138,243 high-quality cells.

In the multiome sample, a stem cell cluster in the transcriptomic data was classified as gonadotrophs in the chromatin data (**Figure 2K**), indicating potentially misaligned annotations. Model uncertainty analysis, however, pointed to a potential transitory state in these cells, which specifically expressed *Lef1* (**Figure 2L**), a gene recently linked to stem cell commitment towards the gonadotroph lineage.^33^ This supports the view that chromatin changes precede transcriptional activation, and highlights model uncertainty as a source of biological insight.

In summary, the CTM accurately identifies cell types in unseen data, providing a consistent framework for cell typing. Both CTM and DM can be run independently or together in a single line of code using the epitome_tools Python package (see Methods).

### Pituitary cell types exhibit sex-biased gene expression in response to sex hormones

Previous single-cell profiling studies of the pituitary largely overlooked sex differences^37^ and the only study addressing them used bulk sequencing, which was confounded by cell type composition, identifying 75 sex-biased genes.^59^ To fill this knowledge gap, we compared male and female CPA samples and identified 6,786 instances of sex-biased expression across cell types (**Figure 3A**). The largest differences were observed in immune cells, lactotrophs, gonadotrophs, with sex-bias also present in stem cells (**Figure 3A**).

**Figure 3.**
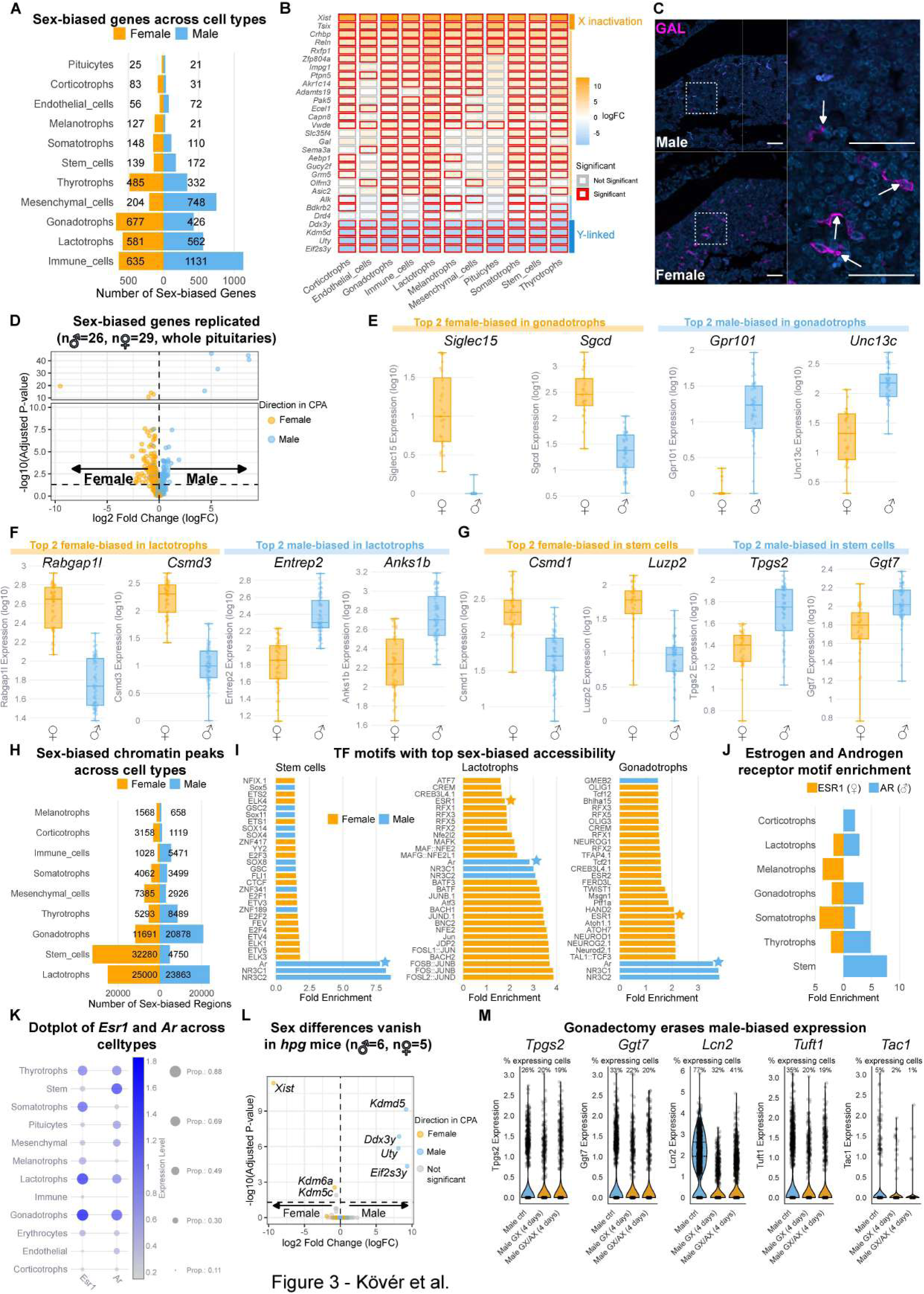
Pituitary cell types exhibit significant sex-bias in gene expression, directed by the activity of Androgen receptor and Estrogen receptor 1. (A) Bar plot showing numbers of statistically significant sex-biased genes (orange: female-biased; blue: male-biased) across cell types. (B) Heatmap of sex-biased genes (log_2_ fold-changes) for those with sex-bias in ≥5 cell types. Orange: female-biased; blue: male-biased. Red rectangles highlight statistically significant cases. (C) Immunofluorescence staining against GAL (magenta) in a male and female pituitary. Scale bars: 50 μm. (D) Volcano plot of sex-biased genes from reanalysis of whole pituitary male/female bulk RNA-seq samples,^59^ colored according to sex-bias in CPA. (E) Box plot of top statistically significant sex-biased genes in gonadotrophs. Left to right: *Siglec15*, *Sgcd*, *Gpr101,* and *Unc13c*. (F) Box plot of top statistically significant sex-biased genes in lactotrophs. Left to right: *Rabgap1l, Csmd3, Entrep2, Anks1b*. (G) Box plot of top statistically significant sex-biased genes in stem cells. Left to right: *Csmd1*, *Luzp2, Tpgs2* and *Ggt7*. (H) Bar plot of statistically significant sex-biased chromatin accessibility peaks (orange: female-biased; blue: male-biased) across cell types, ordered by the absolute number of sex-biased genes. (I) Bar plot of fold-enrichment for top sex-biased TF hits in stem cells, lactotrophs and gonadotrophs. Orange: female-biased; blue: male-biased. Stars highlight ESR1 and AR. (J) Bar plot of TF enrichment in sex-biased chromatin accessibility peaks for ESR1 and AR (orange: female-biased; blue: male-biased). (K) Dot plot of *Esr1* and *Ar* across pituitary cell types. (L) Volcano plot of sex-biased genes from reanalysis of ^67^. Whole pituitary male/female bulk RNA-seq samples from hypogonadal mice, colored according to sex-bias in CPA. (M) Violin plots showing *Tpgs2, Ggt7, Lcn2, Tuft1* and *Tac1* expression in male control, GX (gonadectomy), and GX/AX (gonadectomy/adrenalectomy) samples. Percentages indicate fraction of expressing cells.

We first sought to identify genes with sex-specific expression across multiple cell types. 88 genes showed significant sex-bias in ≥5 cell types (top 30 shown in **Figure 3B**). These included known sex-linked genes (female-biased: *Xist, Tsix*; male-biased: *Uty, Ddx3y, Eif2s3y, Kdmd5*), as well as genes potentially contributing to sex-specific physiological processes, such as *Crhbp,* encoding the corticotropin-releasing hormone binding protein. Another female-biased gene, termed galanin (*Gal*) encodes a positive regulator of ACTH production in corticotrophs^60,61^ and PRL production in lactotrophs,^62,63^ and is stimulated by estrogen.^62,63^ Immunostaining for GAL on male and female mice at P30 confirmed the higher abundance of GAL protein in female pituitaries compared to male (5.7% vs 1.7% of anterior lobe cells; **Figure 3C**).

To validate CPA findings with an independent approach, we uniformly reprocessed male and female bulk RNA-seq data from whole pituitaries,^59^ gonadotrophs,^64^ and corticotrophs.^65^ Genes significant in both CPA and bulk datasets showed near-complete directional concordance (whole pituitary: 188/208 genes, **Figure 3D**; gonadotrophs: 308/314 genes - **Supplementary Figure 3A**; corticotrophs: 73/75 genes, **Supplementary Figure 3B**). We next examined cell type-specific sex-biased expression, focusing on genes showing bias in no more than three cell types. In gonadotrophs, top hits included *Siglec15, Sgcd, Unc13c* and *Gpr101* (**Figure 3E**), the latter associated with X-linked acrogigantism^66^ and showing male-biased expression (**Supplementary Figure 3A, C**). Additionally, *Fshb* and its regulators *Grem1* and *Gata2* were strongly upregulated in male gonadotrophs, confirming previous findings (**Supplementary Figure 3D**).^20^ In lactotrophs, top hits included *Rabgap1l*, *Csmd3*, *Entrep2* and *Anks1b* (**Figure 3F**), while stem cell-specific sex-biased genes included *Csmd1*, *Luzp2, Tpgs2* and *Ggt7* amongst others (**Figure 3G**).

To explore mechanisms underlying sex-biased gene expression, we examined chromatin accessibility patterns. The number of sex-biased peaks across cell types were similar to RNA-level findings, with lactotrophs showing the strongest bias (**Figure 3H**). TF motif analysis revealed strong enrichment for motifs of androgen receptor (AR) and other nuclear receptor motifs (NR3C1 and NR3C2) in male-specific peaks, and estrogen receptor 1 (ESR1) motifs in female-specific peaks (**Figure 3I**). This pattern was consistent across most pituitary cell types (**Figure 3J**). Stem cells lacked ESR1 motif enrichment and showed minimal *Esr1* expression (**Figure 3K**). At the RNA level, none of the cell types exhibit sex bias in *Ar* or *Esr1* (**Supplementary Table 4**), underscoring the added value of chromatin accessibility analysis.

To validate if sex hormones direct expression differences in the pituitary, we uniformly reprocessed a bulk RNA-seq dataset on hypogonadal (*hpg*) mice,^67^ lacking sex hormone production. In this dataset, significant sex differences vanished (**Figure 3L**), except those related to sex chromosomes (female-biased: *Xist, Kdm6a, Kdm5c*; male-biased: *Uty, Ddx3y, Eif2s3y, Kdmd5*). We also retrieved CPA data on stem cells from male controls, male gonadectomized (GX) mice and males following both adrenalectomy (AX) and GX.^22^ Male-biased stem cell genes generally (including top 5: *Tpgs2, Ggt7, Lcn2, Tuft1* and *Tac1;* **Figure 3M**) decreased in GX (**Supplementary Figure 3E**), further supporting the role of sex hormones in driving sex-biased expression.

### Pituitary stem cells show an age-dependent inflammatory gene expression programme

We next explored age-related trends for each cell type, revealing 12,260 instances of age-dependent gene expression (**Supplementary Table 5**; **Figure 4A**). DEGs changing in 3 or more cell types with age, reveal a pituitary-wide increase in inflammation, decrease in cell cycle activity, and a changing WNT-signaling landscape (**Figure 4B**). To confirm these results we interrogated a uniformly reprocessed bulk RNA-seq datasets from whole pituitaries of young and aged mice.^17^ There was strong directional agreement between significant genes in both datasets (201/228 showing identical expression trends **Figure 4C**).

**Figure 4.**
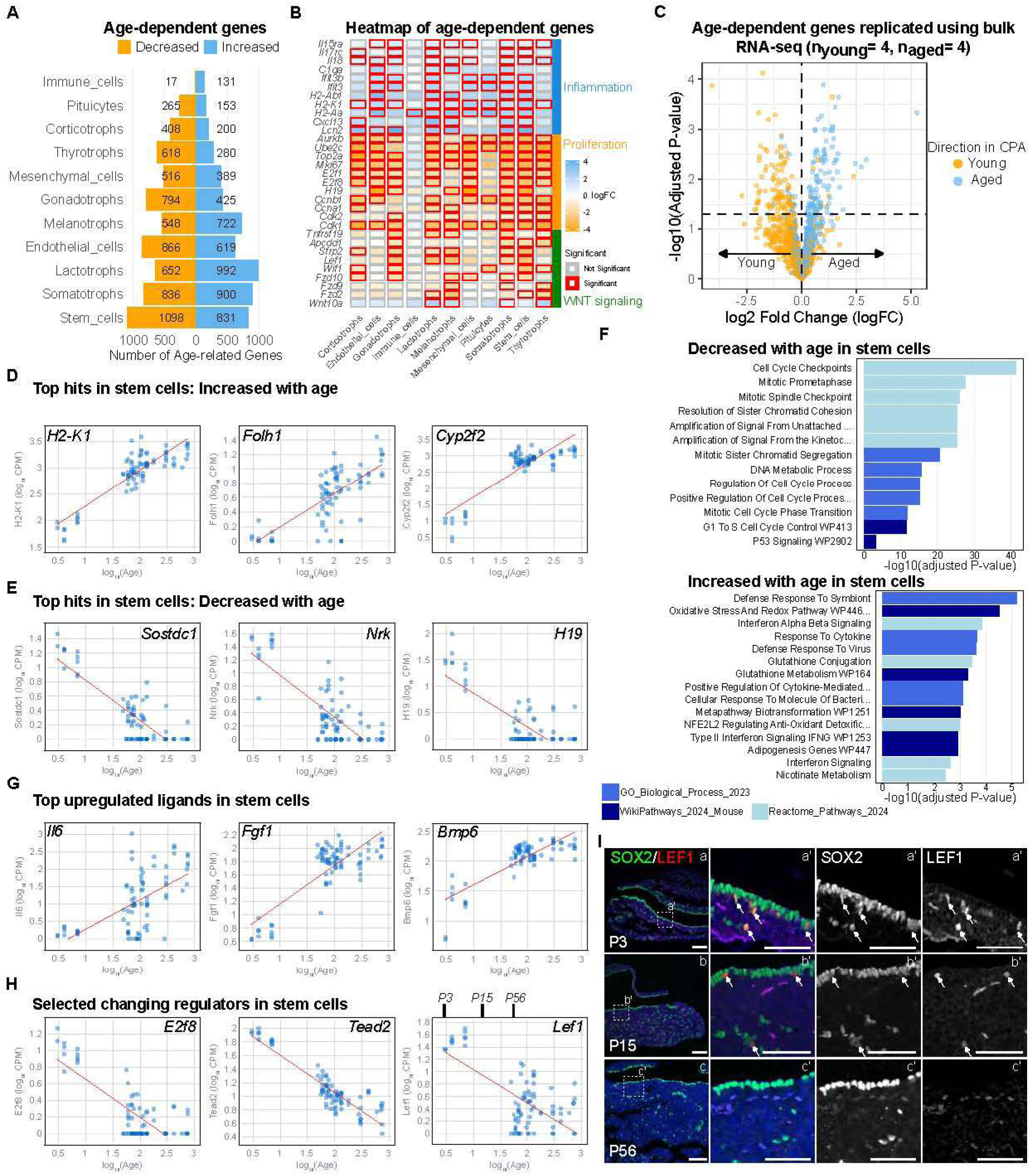
Stem cells exhibit increased inflammation and decreased cell cycle activity with age. (A) Bar plot of statistically significant age-dependent genes (orange: higher in young, blue: higher in aged) across cell types. (B) Heatmap of age-dependent genes (log_2_ fold-changes) for a curated set of genes that show age-dependency in ≥3 cell types. Orange: higher in young, blue: higher in aged. Red rectangles highlight statistically significant cases. (C) Volcano plot from reanalysis of young vs aged whole pituitary RNA-seq samples,^17^ colored according to age-dependency in CPA. (D) Gene expression versus age (log_10_ days) of top 3 statistically significant increasing hits in stem cells. Line of best fit in red. (E) Gene expression versus age (log_10_ days) of top 3 statistically significant decreasing hits in stem cells. Line of best fit in red. (F) Bar plot of enriched terms with genes that increase (top) or decrease (bottom) with age in pituitary stem cells. (G) Gene expression versus age (log_10_ days) of top 3 statistically significant increasing signaling genes in stem cells. Line of best fit in red. (H) Gene expression versus age (log_10_ days) of 3 statistically significant selected TF genes in stem cells. Line of best fit in red. Postnatal ages indicated on *Lef1* plot. (I) Immunofluorescence staining against LEF1 (red) and SOX2 (green) in pituitaries of mice at P3, P15 and P56. Dashed squares a’-c’ are magnified. Scale bars in a-c 100 μm, in a’-c’ 50 μm.

Stem cells exhibited the highest number of age-dependent DEGs (1929), nearly 3 times as many than a previous study (724 DEGs).^37^ The top 3 genes were *H2-K1, Folh1, Cyp2f2* (increasing, **Figure 4D**), and *Sostdc1, Nrk, H19* respectively (decreasing, **Figure 4E**). Enrichment analysis of stem cell-specific age-dependent genes highlighted terms associated with proliferation/cell cycle activity (decreasing) (**Figure 4F**) and inflammatory response and chemokine/cytokine signaling (increasing). Among increasing hits (**Supplementary Table 5**), were genes related to innate immunity (e.g., *Lcn2*, *Ifit3*, *Ifit3b*), multiple cytokines (e.g., *Il6, Il18, Cxcl1, Cxcl13, Ccl24*), MHC-I/II subunits (e.g., *H2-K1*, *H2-D1, H2-Q6, H2-Q7, H2-Ab1, H2-Aa*), complement-system genes (e.g., *C3, C4b*), and interferon-inducible genes (e.g., *ifi47, ifi35, ifi203*). Amongst upregulated ligands in stem cells, we found *Il6 (***Figure 4G***), Il18, Cxcl1, Cxcl13,* and *Ccl24*, all related to inflammation (**Supplementary Table 5**). Notably, intestinal stem cells also upregulate similar cytokines and MHC-II components with age.^68^ Interestingly, aged pituitaries also had an increased proportion of immune cells (**Supplementary Figure 4A**).

The data also revealed a shift in the production of paracrine factors. For example, *Fgf16*, *Bmp2*, *Bmp4*, *Wnt5b*, and *Wnt6* showed downregulation, while *Fgf1, Fgf2, Bmp6, Il6, Il18, Ccl28* and *C3* were upregulated (**Figure 4G, Supplementary Figure 4B**). The upregulated secretory profile mirrors a senescent-associated secretory phenotype (SASP), associated with certain pituitary tumors,^4,69,70^ with several factors included in the SENMAYO gene signature^71^ (P-value: 5.5e-7, Fisher’s exact test). *Cd274*, encoding PD-L1, a well-known immune checkpoint, expressed in pituitary neuroendocrine tumors,^72,73^ was also upregulated (**Supplementary Figure 4B**).

To understand the drivers of these changes, we queried TFs with age-dependent expression (**Supplementary Figure 4C**). Amongst those decreasing, we found cell-cycle regulators *E2f1*, *E2f7*, and *E2f8*, the Hippo pathway effector *Tead2* and WNT/Beta-catenin effector *Lef1* (**Figure 4H**), all of which have the potential to contribute to decreased stem cell contribution with age. Immunofluorescence staining identified LEF1+/SOX2+ cells at P3, and this population decreased by P15 and was undetectable at P56 (**Figure 4I, Supplementary Figure 4D and E**). Amongst increasing TFs, we identified *Nr3c2* (encoding the mineralocorticoid receptor) as well as several others with immune-related functions, such as *Sp100*, *Bhlhe41*, *Stat1*, *Runx1*, and *Tbx21* (encoding T-BET) (**Supplementary Figure 4C**).^74–78^ These TFs might drive the upregulation of cytokines and immune-related genes discussed above.

### Consensus lineage markers of stem cells reveal novel low-expression markers

Differential expression is typically calculated across all cell types present, and therefore has limited biological interpretability for specific lineages. To address this, we derived pituitary “lineage markers” by performing differential expression using cell type comparisons corresponding to branch points in the pituitary lineage (Methods; **Supplementary Table 6**). Comparing stem cells with all committed cells revealed 2307 genes associated with stemness, and a further 1229 genes that are lower in stem cells compared to committed cells. Similarly, we carried out these comparisons for all branch points in the differentiation pathways (**Figure 5A**).

**Figure 5.**
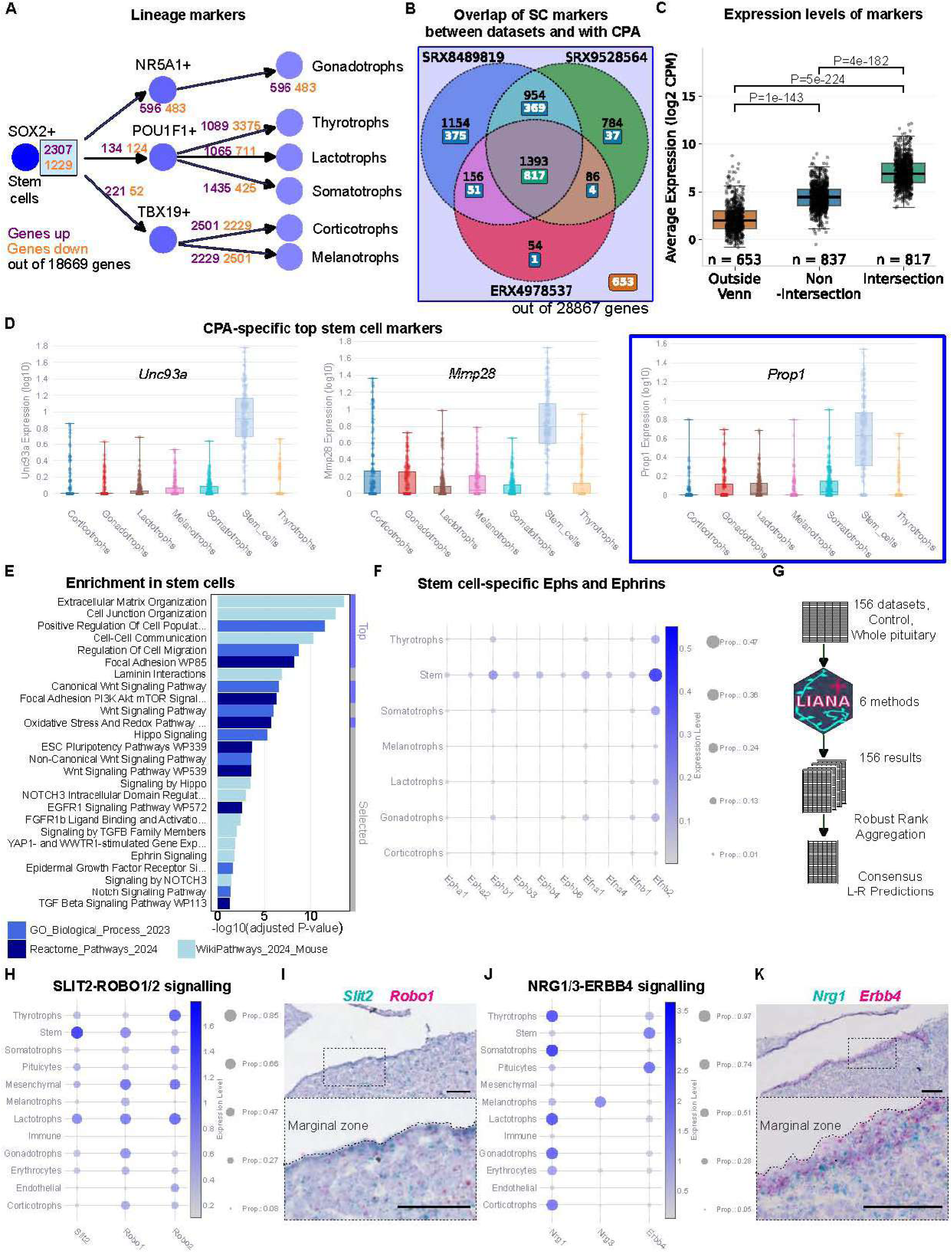
Improved lineage marker detection and transcriptomic analysis using the CPA. (A) Schematic listing the number of lineage markers during anterior pituitary differentiation hierarchy (upregulated: purple, downregulated: orange). (B) Venn diagram of stem cell markers from reanalysis of ^9,13,15^. Boxed numbers indicate overlapping markers derived from the CPA. (C) Box plot of expression values for the three groups of CPA markers from Figure 5B. (D) Box plots of expression values of three low expression markers genes: (Top 2) *Unc93a, Mmp28* and (selected) *Prop1*. (E) Bar plot of top and selected enriched terms in stem cells using three different databases. (F) Dot plot of Ephs and Ephrins enriched in stem cells. (G) Schematic of ligand-receptor interaction prediction. LIANA+ was run on 156 wild-type datasets, followed by robust rank aggregation for a single consensus set of predictions. (H) Dot plot of *Slit2*, *Robo1* and *Robo2* in pituitary cell types. (I) RNAscope mRNA *in situ* hybridization for *Slit2* (red) and *Robo1* (cyan). Scale bars 50 μm and 25 μm (inset). (J) Dot plot of *Nrg1*, *Nrg3* and *Erbb4* in pituitary cell types. (K) RNAscope mRNA *in situ* hybridization for *Nrg1* (cyan) and *Erbb4* (red). Scale bars 50 μm and 25 μm (inset).

To determine if these atlas-scale markers provide additional insights compared to interrogating individual datasets, we reanalyzed three high-quality single-cell datasets (**Figure 5B**).^9,13,15^ There was a great discrepancy between identified stem cell markers, with 1154, 784, and 54 dataset-specific markers (of all markers, 1654 overlapped with CPA). 576 out of 1393 markers consistent across these three datasets (see middle intersection) were not replicable in the CPA (**Figure 5B; Supplementary Figure 5A**). We attribute these inconsistencies to the statistical limitations of working with a single dataset, as well as the sample-to-sample variation between animals and labs. Furthermore, the CPA showed a particular advantage over these individual samples in capturing an additional 653 lowly-expressed markers (**Figure 5C**; Top 2 hits were *Unc93a* and *Mmp28* for stem cells; **Figure 5D**), that would not have been identified using either single dataset. Amongst these low-expression markers was *Prop1* (**Figure 5D**), a TF with a well-established role in the commitment of pituitary stem cells^79,80^ that has been problematic to detect in single-cell studies.^8^ These results are in line with pseudobulk analysis being more sensitive to low-expression genes.^53^ We then also identified additional markers (including low-expression ones) compared to individual datasets, for all other points in the pituitary lineage (**Supplementary Figure 5B; Supplementary Table 7**). Strikingly, from the 2501 upregulated genes in corticotrophs compared to melanotrophs in the CPA, only 52 would have been uncovered by the intersection of the 3 individual datasets (**Supplementary Figure 5B**). Overall, these results demonstrate the power of using the CPA for marker discovery compared to individual datasets.

We then performed gene set enrichment on stem cell lineage markers, which revealed known pituitary stem cell pathways (**Supplementary Table 8**), such as WNT-signaling,^81^ EGF signaling, FGF signaling, Notch signaling,^82,83^ and the Hippo pathway^1,84,85^ (**Figure 5E**). Additional enriched terms included Eph-Ephrin signaling and Laminin interactions (**Figure 5E**). Specifically, several low-expression Ephrins (*Efna1, Efna4, Efnb1, Efnb2*) and Eph receptors (*Epha1, Epha2, Ephb1, Ephb3, Ephb4, Ephb6*) are specific to stem cells (**Figure 5F**), while other Ephs and Ephrins are expressed across pituitary cell types (**Supplementary Figure 5C**). The laminin genes enriched in stem cells included *Lama3 Lama5, Lamb2, Lamb3*, *Lamc1* and *Lamc2* (**Supplementary Figure 5D**).

### Consensus ligand-receptor interactions reveal paracrine signaling between stem and endocrine cells

Since stem cell lineage markers were enriched for cell-cell communication genes (**Figure 5E**), we next investigated ligand-receptor interactions across all control, whole-pituitary sc/snRNA-seq datasets (156 samples, 598,993 cells). We applied LIANA+,^86^ which integrates multiple ligand-receptor inference algorithms, and merged results using robust rank aggregation^87^ (Methods; **Figure 5G; Supplementary Table 9**). Cell type-specific interactions are summarized in **Supplementary Figure 5E**.

For instance, stem cells showed high *Slit2* expression encoding SLIT2, predicted to signal through ROBO1 and ROBO2 on differentiated cell types (**Figure 5H**). RNAscope mRNA *in situ* hybridization confirmed this pattern showing *Slit2* localization near the marginal zone (stem cell niche) and *Robo1* in adjacent cells (**Figure 5I**).

Additional predicted stem cell interactions included the ERBB4 receptor engaging with NRG3 from melanotrophs and NRG1 from other differentiated cell types (**Figure 5J**). RNAscope mRNA in situ hybridization confirmed *Erbb4* expression near the marginal zone and *Nrg1* in adjacent cells (**Figure 5K**). Stem cells were also predicted to be the sole source of BMP6, while all other cell types variably expressed its receptors (*Bmpr1a*, *Bmpr2a*, *Acvr1* and *Acvr2a*) (**Supplementary Figure 5E, F**). In addition, stem cells emerged as the main source of *Fgf1* expression along with corticotrophs; receptor genes *Fgfr1* and *Fgfr2* were expressed primarily by stem cells and broadly across other cell types, respectively (**Supplementary Figure 5E, G**). Finally, this approach also highlighted TGFB2-TGFBR1/2/3 interactions, with *Tgfb2* expression restricted to stem cells and receptor expression detected across all cell types (**Supplementary Figure 5E, H**).

These results revealed several highly-specific interactions between all cell types, including many between pituitary stem cells and endocrine cells, reinforcing the notion of pituitary stem cells as paracrine signaling hubs.^88,89^

### Novel candidate TFs associated with cell fates across modalities

Building on gene expression analyses, we examined chromatin accessibility changes along the pituitary lineage (**Supplementary Table 10**). This identified 30,214 peaks enriched in stem cells, and 28,697 peaks more accessible in committed lineages (**Figure 6A**). Multimodal hits were then defined at each cell fate decision point (**Figure 6B and 6C; Supplementary Table 11**), based on TFs that were both differentially expressed and whose motifs were enriched in differentially accessible peaks. Because this approach depends on known binding motifs and also excludes TFs whose accessibility remains unchanged, we also generated an “RNA-only” list (**Supplementary Figure 6A and 7B**; **Supplementary Table 11**).

**Figure 6.**
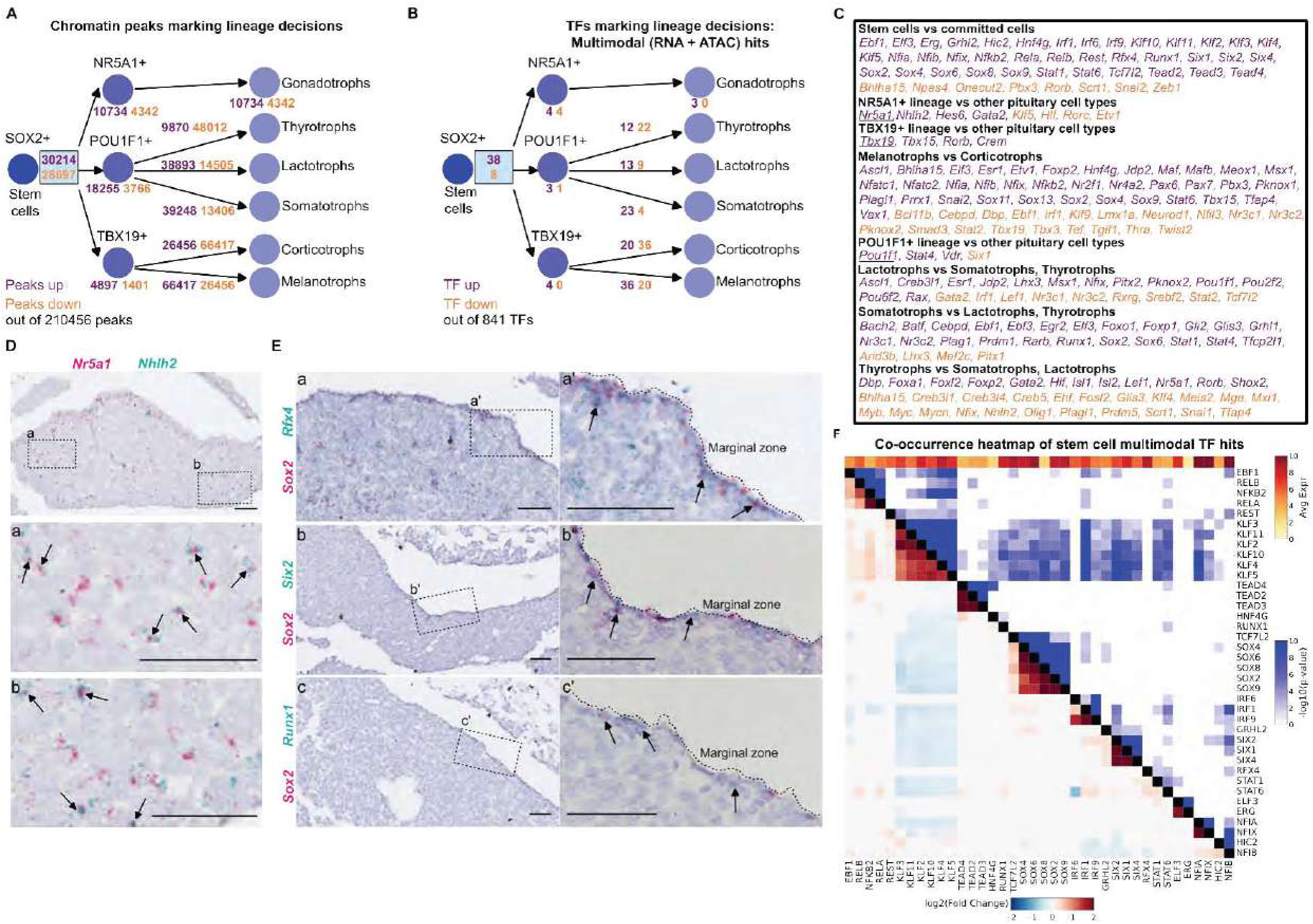
Identification of fate-specific transcription factors through transcriptomics and chromatin accessibility. (A) Schematic of the number of lineage marker peaks in the differentiation hierarchy of the pituitary lineage (opening: purple, closing: orange). (B) Schematic of the number of Multimodal hit TFs in the differentiation hierarchy of the pituitary lineage (upregulated: purple, downregulated: orange). (C) Table of all Multimodal hits. (upregulated: purple, downregulated: orange). (D) RNAscope mRNA *in situ* hybridization for *Nr5a1* (red), and *Nhlh2* (cyan). Scale bar 50 μm. (E) RNAscope mRNA *in situ* hybridization for target TFs. Top: *Sox2* (red), *Rfx4* (cyan); Middle: *Sox2* (red), *Six2*: cyan; Bottom: *Sox2* (red), *Runx1* (cyan). Scale bar 50 μm. (F) Motif co-occurrence heatmap of Multimodal hit TFs in stem cells. Rows and columns share the same TF order. Colors below the diagonal represent fold change of observed versus expected co-occurrence rates, while colors above indicate statistical significance (–log₁₀ p-value).

The TBX19+ lineage contained 4 Multimodal hits: *Tbx19*, *Tbx15*, *Rorb and Crem* (**Figure 6B and 6C**). RNA-only hits included *Esrrb* and *Zim3* (**Supplementary Figure 6B**; **Supplementary Table 11**). Within this lineage, several Multimodal hits differed between corticotrophs and melanotrophs. Among these, the melanotroph TF-encoding gene *Pax7* is well-established as a determinant of melatotroph over corticotroph fate,^90^ while *Etv1* encodes a key transcriptional regulator active in this lineage.^91^ Corticotroph-specific TFs included *Neurod1* and the glucocorticoid receptor *Nr3c1*. *Neurod1* is required for proper corticotroph differentiation,^92^ regulates *Pomc* transcription^93,94^ and is expressed in corticotroph adenomas.^95^

In the NR5A1+ lineage, we identified *Nhlh2, Gata2* and *Hes6* as additional Multimodal hits (**Figure 6B and 6C**). To confirm the specificity of *Nhlh2* in gonadotrophs, we performed double RNAscope mRNA *in situ* hybridization for *Nhlh2* and *Nr5a1*, which showed clear co-expression (**Figure 6D**). In gonadotrophs, *Nhlh2* displayed female-biased expression (**Supplementary Figure 6C**), of relevance for future functional studies. RNA-only hits included *Foxl2* and *Foxp2* amongst others (**Supplementary Figure 6B**; **Supplementary Table 11**).

In addition to *Pou1f1*, the POU1F1+ lineage contained 2 additional Multimodal hits (**Figure 6B and 6C**); *Stat4* and the vitamin D receptor *Vdr*, whose homozygous mutants show reduced growth and IGF-1 levels.^96,97^ RNA-only hits included the retinoid X receptor gamma *Rxrg*, involved in growth regulation and thyrotroph function^28,98,99^ (**Supplementary Figure 6B**; **Supplementary Table 11**). RXRG and VDR can act cooperatively, and have previously been implicated in *Pou1f1* regulation.^100–102^

Within the POU1F1+ lineage, we also resolved TF-encoding genes that might drive differentiated cell type identities. Somatotroph-specific hits included genes previously implicated in somatotroph differentiation *Gli2*,^103,104^ *Foxo1,*^105,106^ and *Nr3c1.*^107,108^ Thyrotroph-specific hits interestingly included *Nr5a1*, despite its canonical role in gonadotroph fate, as well as various other previously identified genes, such as *Gata2, Isl1, Foxl2*, and *Shox2*.^23^ Lactotroph Multimodal hits included *Esr1* encoding estrogen receptor, as well as *Pou6f2*. Interestingly, *POU6F2* mutations have been identified in prolactinomas.^109^

Multimodal hits associated with stem cells included genes from the SOX (*Sox2, Sox4, Sox6, Sox8, Sox9*) and TEAD (*Tead2, Tead3, Tead4*) families, both known regulators of pituitary stem cells, were identified.^1,4,85^ Additional TF families included KLF (*Klf2, Klf3, Klf4, Klf5, Klf10, Klf11*), NFI (*Nfia, Nfib, Nfix*), SIX (*Six1, Six2, Six4*), IRF (*Irf1, Irf6, Irf9*), STAT (*Stat1, Stat6*), REL (*Rela, Relb*), along with single TF-encoding genes (*Rfx4, Runx1, Grhl2, Rest, Nfkb2, Hic2, Hnf4g, Erg, Tcf7l2, Elf3* and *Ebf1*). Members of the TEAD, NFI and SIX families showed the strongest enrichment in stem cell-associated chromatin regions (**Supplementary Figure 6D**). Double RNAscope mRNA *in situ* hybridization confirmed *Runx1, Rfx4* and *Six2* expression alongside *Sox2* in pituitary stem cells (**Figure 6E**).

We then examined motif co-occurrence across stem cell-specific Multimodal hits (**Figure 6E**). Interestingly, KLF binding sites were less likely than expected to co-occur with other TF motifs within the same peaks (**Figure 6F**). Genomic annotation of KLF motifs showed that they were enriched in proximal regulatory regions relative to other TF motifs (**Supplementary Figure 6E and 7F**), suggesting a functional basis for their limited overlap.

### The *epitome*: a programming-free interface for the Consensus Pituitary Atlas

Democratized access to the CPA is key to advancing reproducibility and collaboration. To this end, we developed the electronic pituitary omics (*epitome*) platform,^49^ which currently hosts the CPA at epitome-atlas.com and will be continuously updated with published datasets. Through *epitome*, users can perform extensive exploratory analyses online across published transcriptomic and chromatin accessibility datasets. Available secondary analyses include LIANA+ ligand-receptor interactions,^86^ gene-gene coexpression, age-related expression trends, TF-binding motif visualization, TF-binding motif enrichment via ChromVAR,^110^ and generating TF motif co-occurrence heatmaps. The platform generates publication-ready figures for all these analyses, facilitating seamless integration into research workflows. Uniformly processed datasets are also available for download, enabling local analysis, as well as automated processing and annotation (through Cell Type and Doublet models) of users’ own data, all without programming or CPA data submission.

## Discussion

Fulfilling their promise of cell type-specific insights, single-cell genomic methodologies are truly effective when carried out at a grand scale. Here we present the Consensus Pituitary Atlas, a comprehensive and scalable resource, encompassing all currently available mouse pituitary single-cell datasets. The CPA enables statistically robust analyses and serves as a definitive reference for this organ. As proof-of-concept, we use the CPA to reveal sex- and age-related effects on gene regulatory programmes at unprecedented resolution, underscoring the power of a domain-specific atlas that can be readily extended to other tissues and species.

The CPA also enables more accurate marker identification than individual studies, yielding numerous robust cell type-specific TF candidates, including previously undescribed regulators. For example, *Rfx4* emerges as a stem cell TF, showing stem cell-specific expression and chromatin accessibility. Pathogenic *RFX4* variants have been reported in six patients with neurodevelopmental disorders, two of whom also presented pituitary anomalies including agenesis,^111^ supporting its functional relevance. Cell-cell communication analyses identified SLIT-ROBO signaling among top pathways. Multiple hypopituitarism patients carry *ROBO1* variants,^112–116^ previously linked to defects in the neural pituitary.^113^ Our finding that stem cells and lactotrophs may secrete SLIT2, signaling via ROBO1/2 receptors across endocrine cell types, offers new insights into anterior pituitary phenotypes. Thus, a CPA framework can guide prioritization for future pre-clinical functional studies.

Analysis of Multimodal hits at lineage decision points, successfully recovered the three key TFs of commitment (NR5A1, POU1F1 and TBX19) and revealed additional candidates whose roles remain unexplored in pituitary differentiation. For instance, *Nhlh2* was an additional TF gene identified alongside *Nr5a1* in gonadotrophs. Although *Nhlh2^−/−^*knockout mice retain a gonadotroph population, they exhibit hypogonadism^117^ so far linked to hypothalamic kisspeptin neurons,^118^ and also display pituitary hypoplasia and downregulated *Gnrhr* expression in gonadotrophs,^119^ suggesting a direct role in pituitary development.

The CPA also provides a framework to study biological ageing and sexual dimorphism. For example, *Crhbp*, encoding corticotropin-releasing hormone binding protein, showed strong female bias across all cell types, being nearly absent in males. As CRHBP sequesters corticotropin-releasing hormone, a key hypothalamic regulator of ACTH, this difference may underlie sex-biased stress responses. Indeed, *Crhbp^−/−^* knockout phenotypes exhibit sex-specific gene expression and behavioral patterns relating to stress.^120,121^ The CPA also uncovered an age-associated inflammatory programme within pituitary stem cells, revealing coordinated transcriptional and paracrine shifts reminiscent of tissue senescence. Ageing stem cells showed marked upregulation of cytokines, interferon-responsive and complement genes, together with immune checkpoint molecules such as *Cd274* (PD-L1), while proliferative and stem cell-related factors, including *E2f1*, *Tead2*, and *Lef1* declined. This dual profile, heightened immune-related paracrine signaling and reduced regenerative potential, mirrors a senescence-associated secretory phenotype^71^ and may contribute to functional decline in the pituitary with age. The age-dependent upregulation of immune-related TFs (*Bhlhe41, Sp100, Stat1, Runx1*, *Tbx21*) further points to a transcriptional reprogramming of stem cells toward an inflammatory state, similar to those reported in intestinal stem cells^68^ and in the ageing brain.^122^ BHLHE41 potentially represses the E2F network that participates in proliferation,^78^ whereas STAT1 has been previously shown to drive inflammaging in intestinal stem cells,^68^ upregulating similar genes to those observed here. Together, these findings suggest that pituitary ageing involves an intrinsic, stem cell-centred inflammatory remodeling, which could influence susceptibility to endocrine insufficiency or tumorigenesis later in life.

Finally, the CPA is complemented by the *epitome* platform,^49^ a programming-free, user-friendly interface for exploring and analyzing the atlas, and in line with the FAIR guiding principles.^123^ Epitome allows users to interrogate all existing mouse datasets, or to uniformly process and analyze their own data locally, without prior submission. The accompanying epitome_tools Python package further enables rapid, expert-level cell type annotation and doublet filtering via machine learning, using a single command. Together these resources democratize data access, promote standardized best practices, and accelerate discovery without requiring programming expertise. Looking forward, both the atlas and platform can expand to additional modalities, such as spatial transcriptomics, single-nucleus methylomics and single-cell proteomics.

## Limitations of the study

This work predicts various TF-encoding genes based on gene expression and chromatin accessibility. Chromatin accessibility, highlights enrichment in chromatin peaks with changing accessibility but does not prove TF binding, which requires functional assays. Across our analyses, results for POU1F1+, TBX19+ and NR5A1+ intermediate progenitors are stated; however, these populations are transient, and may not be sufficiently captured by single-cell approaches in postnatal samples. Future datasets could enrich for these transient cells by various cell sorting strategies to directly resolve their intermediate transcriptomic profiles.

## Resource availability

### Lead contact

Further information and requests for resources and reagents should be directed to and will be fulfilled by Prof. Cynthia Andoniadou.

### Materials availability

This study did not generate new unique reagents or organisms.

### Data and code availability

All generated code is available on Github: https://github.com/Andoniadou-Lab/epitome, https://github.com/Andoniadou-Lab/epitome_tools, https://github.com/Andoniadou-Lab/consensus_pituitary_atlas.

The epitome_tools Python package can be installed via pip install epitome_tools and used following the GitHub instructions. Processed files and an integrated RNA/ATAC object are available on the epitome platform (“Downloads” tab). Pseudobulk and single-cell objects, as well as website code, are deposited in Zenodo (10.5281/zenodo.17359014; doi:10.5281/zenodo.17154160). The multiome dataset is deposited in the Sequence Read Archive (PRJNA1329231).

The generated multiome dataset was deposited in the Sequence Read Archive (PRJNA1329231).

## Acknowledgements

Thanks to Ian Kruk for feedback on the *epitome* platform and Zsolt Balla (KCL e-research) for help with the *epitome* website. Sequencing was performed at the New York Genome Center. We thank Scientific Computing at Mount Sinai for computational resources and support.

This work was funded by the Medical Research Council (grants APP40962 and MR/T012153/1) and the Deutsche Forschungsgemeinschaft (DFG, German Research Foundation), Project no. 314061271, TRR 205: “The Adrenal: Central Relay in Health and Disease” and Project no. 288034826, IRTG 2251: “Immunological and Cellular Strategies in Metabolic Disease” to CLA. BK and MVS were funded by the Wellcome Trust as part of the Advanced Therapies for Regenerative Medicine Wellcome Trust PhD Programme (218461/Z/19/Z). TLW was funded by King’s College London as part of the ‘Cell and Therapies and Regenerative Medicine’ Four-Year Wellcome Trust PhD Training Programme (108874/Z/15/Z). DJB was funded by Canadian Institutes of Health Research Project Grants PJT-191766 and PJT-195832. SCS was supported by funding from the National Institutes of Health (NIH) Grant DK46943. FRZ received Cedars-Sinai institutional support.

## Author Contributions

Conceptualization: BK, CLA; Data curation: BK; Formal analysis: BK., OS; Funding acquisition: CLA; Investigation: BK, TLW, OS, JKC, YK, MVS, EL, FRZ, NM, MZ, ZZ, GRS; Methodology: BK, HCL, CLA; Project administration: CLA; Resources: TLW, CLA; Software: BK; Supervision: DJB, HCL, SCS, FRZ, CLA; Visualization: BK; Writing - original draft: BK, CLA; Writing - review & editing: BK, CLA, HCL, FRZ, DJB, SCS, TLW, OS, JKC, YK, MVS, EL, FRZ, NM, MZ, ZZ, GRS.

## STAR Methods

### Key resources table

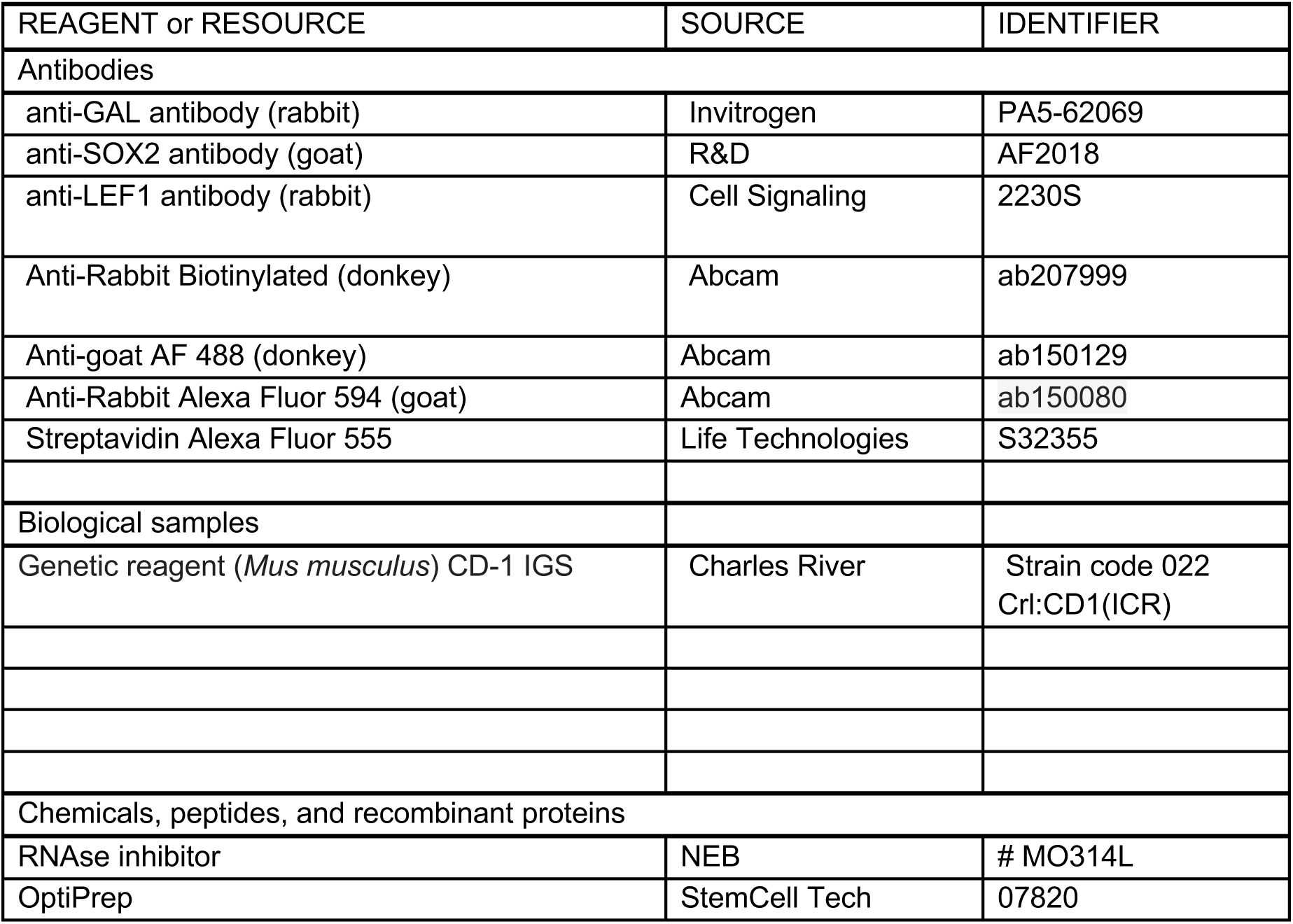
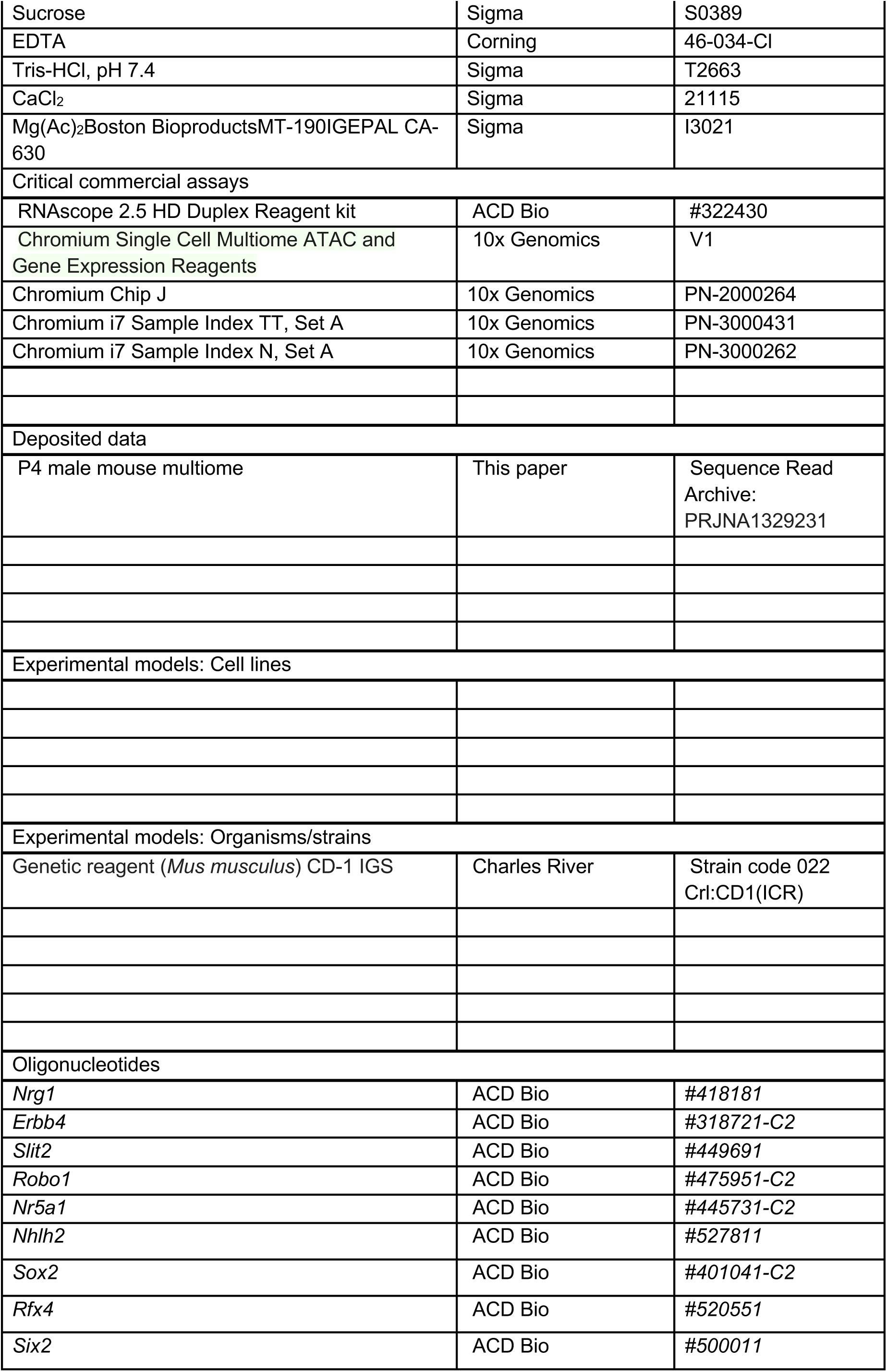
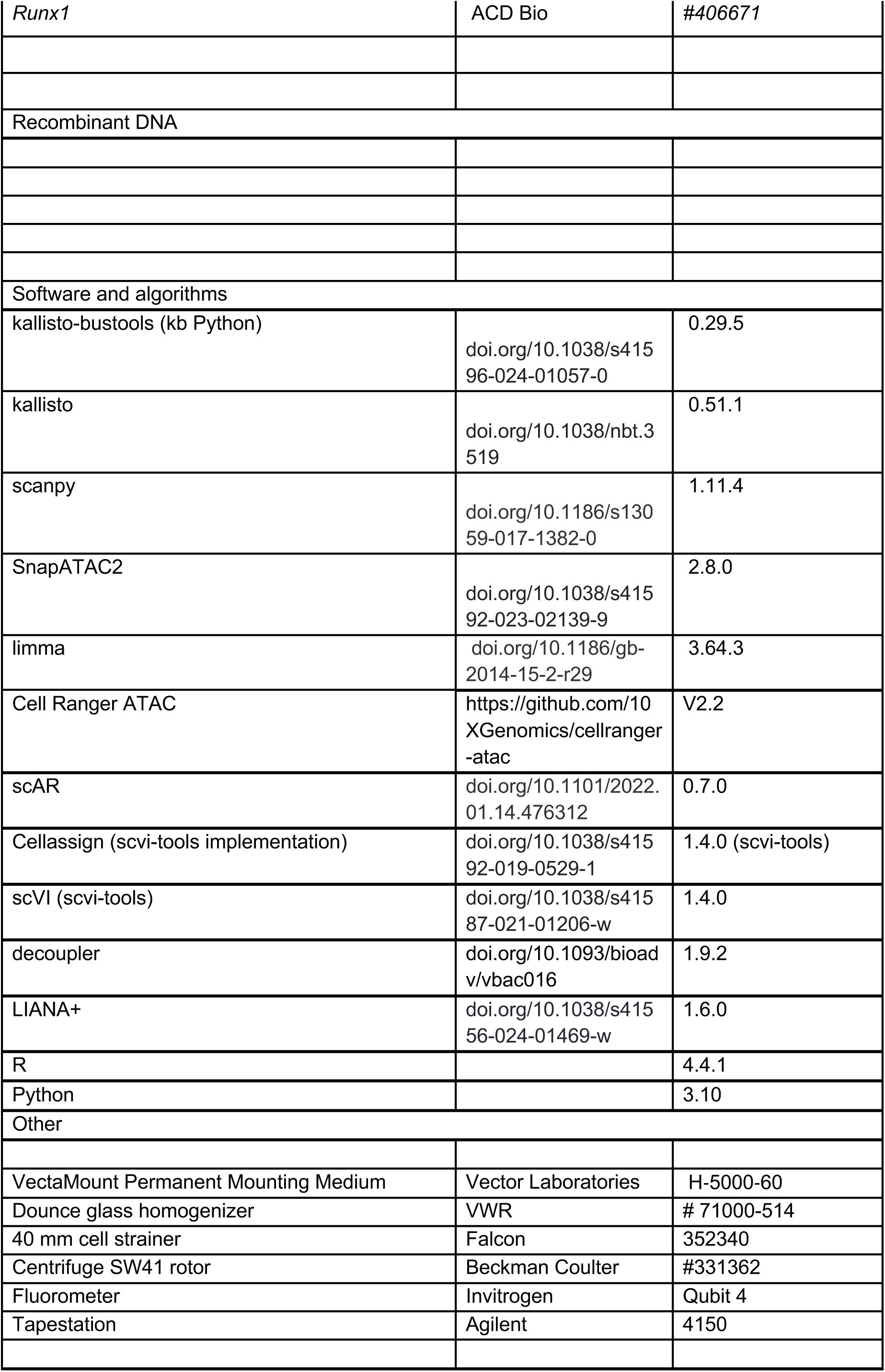

| REAGENT or RESOURCE | SOURCE | IDENTIFIER |
| --- | --- | --- |
| Antibodies |  |  |
| anti-GAL antibody (rabbit) | Invitrogen | PA5-62069 |
| anti-SOX2 antibody (goat) | R&D | AF2018 |
| anti-LEF1 antibody (rabbit) | Cell Signaling | 2230S |
| Anti-Rabbit Biotinylated (donkey) | Abcam | ab207999 |
| Anti-goat AF 488 (donkey) | Abcam | ab150129 |
| Anti-Rabbit Alexa Fluor 594 (goat) | Abcam | ab150080 |
| Streptavidin Alexa Fluor 555 | Life Technologies | S32355 |
| Biological samples |  |  |
| Genetic reagent (*Mus musculus*) CD-1 IGS | Charles River | Strain code 022<br>CrI:CD1(ICR) |
| Chemicals, peptides, and recombinant proteins |  |  |
| RNAse inhibitor | NEB | # MO314L |
| OptiPrep | StemCell Tech | 07820 |
| Sucrose | Sigma | S0389 |
| EDTA | Corning | 46-034-Cl |
| Tris-HCl, pH 7.4 | Sigma | T2663 |
| CaCl<sub>2</sub> | Sigma | 21115 |
| Mg(Ac)<sub>2</sub>Boston BioproductsMT-190IGEPAL CA-630 | Sigma | I3021 |
| Critical commercial assays |  |  |
| RNAscope 2.5 HD Duplex Reagent kit | ACD Bio | #322430 |
| Chromium Single Cell Multiome ATAC and Gene Expression Reagents | 10x Genomics | V1 |
| Chromium Chip J | 10x Genomics | PN-2000264 |
| Chromium i7 Sample Index TT, Set A | 10x Genomics | PN-3000431 |
| Chromium i7 Sample Index N, Set A | 10x Genomics | PN-3000262 |
| Deposited data |  |  |
| P4 male mouse multiome | This paper | Sequence Read Archive:<br>PRJNA1329231 |
| Experimental models: Cell lines |  |  |
| Experimental models: Organisms/strains |  |  |
| Genetic reagent (*Mus musculus*) CD-1 IGS | Charles River | Strain code 022<br>Crl:CD1(ICR) |
| Oligonucleotides |  |  |
| *Nrg1* | ACD Bio | #418181 |
| *ErbB4* | ACD Bio | #318721-C2 |
| *Slit2* | ACD Bio | #449691 |
| *Robo1* | ACD Bio | #475951-C2 |
| *Nr5a1* | ACD Bio | #445731-C2 |
| *Nhlh2* | ACD Bio | #527811 |
| *Sox2* | ACD Bio | #401041-C2 |
| *Rfx4* | ACD Bio | #520551 |
| *Six2* | ACD Bio | #500011 |
| *Runx1* | ACD Bio | #406671 |
| Recombinant DNA |  |  |
| Software and algorithms |  |  |
| kallisto-bustools (kb Python) | doi.org/10.1038/s41596-024-01057-0 | 0.29.5 |
| kallisto | doi.org/10.1038/nbt.3519 | 0.51.1 |
| scanpy | doi.org/10.1186/s13059-017-1382-0 | 1.11.4 |
| SnapATAC2 | doi.org/10.1038/s41592-023-02139-9 | 2.8.0 |
| limma | doi.org/10.1186/gb-2014-15-2-r29 | 3.64.3 |
| Cell Ranger ATAC | https://github.com/10XGenomics/cellranger-atac | V2.2 |
| scAR | doi.org/10.1101/2022.01.14.476312 | 0.7.0 |
| Cellassign (scvi-tools implementation) | doi.org/10.1038/s41592-019-0529-1 | 1.4.0 (scvi-tools) |
| scVI (scvi-tools) | doi.org/10.1038/s41587-021-01206-w | 1.4.0 |
| decoupler | doi.org/10.1093/bioadv/vbac016 | 1.9.2 |
| LIANA+ | doi.org/10.1038/s41556-024-01469-w | 1.6.0 |
| R |  | 4.4.1 |
| Python |  | 3.10 |
| Other |  |  |
| VectaMount Permanent Mounting Medium | Vector Laboratories | H-5000-60 |
| Dounce glass homogenizer | VWR | # 71000-514 |
| 40 mm cell strainer | Falcon | 352340 |
| Centrifuge SW41 rotor | Beckman Coulter | #331362 |
| Fluorometer | Invitrogen | Qubit 4 |
| Tapestation | Agilent | 4150 |

### Experimental model and study participant details

Animal husbandry was carried out under compliance of the Animals (Scientific Procedures) Act 1986, Home Office license and KCL ethical review approval. Pituitaries from CD-1 mice were collected from male and randomly-cycling female mice. Animals were on a 12-hour on, 12-hour off light cycle (lights on at 7 AM; off at 7 PM).

A male mouse with mixed CD-1 and C57BL6/J background (backcrossed on CD-1 for 5 generations) was collected for multiome data generation at postnatal day 4.

A male and female mouse with CD-1 backgrounds aged P30 were used for immunofluorescence staining against GAL, whereas mice with mixed background and ages P3, P15 and P56 were used for SOX2/LEF1 stainings.

Mice with CD-1 backgrounds aged P21 were used for RNAscope *in situ* hybridization targeting *Nrg1* and *Erbb4*.

Mice with CD-1 backgrounds aged P14 were used for RNAscope *in situ* hybridization targeting *Slit2* and *Robo1*.

Mice with CD-1 backgrounds aged P14 were used for RNAscope *in situ* hybridization targeting *Nr5a1* and *Nhlh2*.

Mice with CD-1 backgrounds aged P14 were used for RNAscope *in situ* hybridization targeting *Rfx4-Sox2* and P56 for *Runx1* or *Six2* and *Sox2*.

### Method details

#### Dataset curation

All publications with mouse pituitary gland data have been identified through searching Pubmed, GEO or SRA using “pituitary single-cell” or “pituitary single-nucleus” terms. Associated publications were not identified for three datasets (GSE242296^32^, GSE299835^42^, GSE239316^43^) and metadata was exclusively extracted for GEO in these cases. All respective SRA, ENA or GEO identifiers were extracted manually, and organized in a table with available metadata, including genotype, age, sex, estrous cycle stage, assay modality, and single-cell technology (**Supplementary Table 1**). Barcoding kits were confirmed manually by comparing a sample of R1 reads against all 10X Genomics barcode whitelists. This search revealed multiple published datasets with incorrectly stated barcoding kits in the Methods section of the respective publication, its GEO site, or in some cases both. This approach for identifying the correct barcoding chemistry is similar to that in ^47^, where the authors have also noted that barcoding chemistry is unreliable in SRA/GEO metadata.

#### Accessing transcriptomics datasets

Using the SRA IDs, raw sequencing files for datasets^8–43^ were obtained from the Sequence Read Archive (SRA) using ffq^124^ and sra-tools, or manually from ArrayExpress. In some cases, we only found the R2 reads uploaded,^8,28^ which then required us to access the .bam files and convert them to fastq format using the 10X bamtofastq software. The data accession workflow that requires a curated data frame as input is implemented in the `atlas` module of the epitome_tools Python package to enable future atlasing efforts.

#### Accessing IGVF samples

The samples in the IGVF resource have been produced using an elaborate design of splitting and pooling, also mixing various experimental animals which were then deconvoluted using their known genotypic differences (see Rebboah et al.^41^). Recovering information on cells from these samples requires reverse-engineering the split-pool design, as well as mapping to the different strain-specific reference transcriptomes. Given that the alignment software was the same as used in our case, we decided to retrieve the processed AnnData objects (not raw .fastq files) from the IGVF data portal (https://data.igvf.org/analysis-sets/IGVFDS1666GZSV/). Here the CellBender counts layer was used, and cells were subsetted to include pituitary cell types (as annotated by the authors). Following this, the samples were ran through the rest of the QC workflow, except removal of ambient RNA (as it has been done using CellBender already by the authors). We then performed cell typing in accordance with the rest of the samples.

#### Pre-processing - Single-cell transcriptomics

All resulting fastq files were then uniformly pre-processed using the kallisto-bustools v0.51 program^125,126^ using a workflow that quantifies nascent and coding transcripts (“nac”), which captures intronic, exonic and ambiguous reads. The pseudoalignment was against the default reference transcriptome in the command “kb ref” (ensembl release 108 - GRCm39). The final count matrices for each dataset contained a sum of intronic, exonic, and ambiguous matrices. This approach is both up-to-date with the current default in Cell Ranger, and also enables a more uniform quantification across assays (previously intronic regions would only be included in single-nucleus data, but not for single-cell) as highlighted in ^47^.

#### Nuclei isolation from pituitary

The flash-frozen mouse pituitary was processed as an individual sample. Nuclei isolation was performed based on a modified protocol from ^127^, refer to ^128, 129^. Briefly, and all on ice, RNAse inhibitor (NEB cat# MO314L) was added to the homogenization buffer (0.32 M sucrose, 1 mM EDTA, 10 mM Tris-HCl, pH 7.4, 5mM CaCl_2_, 3mM Mg(Ac)_2_, 0.1% IGEPAL CA-630), 50% OptiPrep (Stock is 60% Media from StemCell cat# 07820), 35% OptiPrep and 30% OptiPrep right before isolation. Each sample (single or pooled pituitaries) was homogenized in a Dounce glass homogenizer (1ml, VWR cat# 71000-514), and the homogenate was filtered through a 40 μm cell strainer. An equal volume of 50% OptiPrep was added, and the gradient was centrifuged (SW41 rotor at 17,792xg; 4 °C; 25min). Nuclei were collected from the interphase, washed, resuspended in 1X nuclei dilution buffer, and counted using a fluorescent assay on a K2 Cellometer (Revvity).

#### Sn multiome assay

Sn multiome was performed following the Chromium Single Cell Multiome ATAC and Gene Expression Reagent Kits V1 User Guide (10x Genomics, Pleasanton, CA). Nuclei were counted using propidium iodide fluorescence (Revvity Cellometer counter), transposition was performed in 10 μl at 37 °C for 60 min, targeting 500-20,000 nuclei, before loading of the Chromium Chip J (PN-2000264) for GEM generation and barcoding. Following post-GEM cleanup, libraries were pre-amplified by PCR, after which the sample was split into three parts: one part for generating the snRNA-seq library, one part for the snATACseq library, and the rest was kept at −20C. SnATAC and snRNA libraries were indexed for multiplexing (Chromium i7 Sample Index N, Set A kit PN-3000262, and Chromium i7 Sample Index TT, Set A kit PN-3000431 respectively).

#### Quality control (QC) and sequencing of sn libraries

Libraries were quantified by Qubit 3 fluorometer (Invitrogen), and quality was assessed by Bioanalyzer (Agilent). Equivalent molar concentrations of libraries were pooled, and the reads were adjusted after sequencing the pools in a MiSeq (Illumina). The libraries were then sequenced on a NovaSeq 6000 (Illumina) at the New York Genome Center (NYGC) following recommendations from Illumina and 10X Genomics. Refer to ^128^ for details.

#### Processing Multiome samples

In accordance with the rest of the atlas, the RNA part of the generated multiome samples were aligned using kallisto-bustools and further processing and QC was done as for the other samples.

The ATAC part of the datasets was aligned using Cell Ranger ATAC 2.0, and the rest of the processing and QC was the same as described for the other samples.

#### Further processing and quality control of all sc/snRNA-seq samples

To separate true cells from cell-free droplets, we first filtered to the top 40000 barcodes, which we empirically found close to the knee of the knee plot (log_UMIs vs log_rank plot), given that all datasets ranged between 500 - 20000 cells. Following this, we used the filter command from the mx package,^48^ which fitted two Gaussians on the data to separate true cells and contaminated droplets. In some cases, mx filter gave too few cells (<2000), often with a very high minimum number of UMIs (e.g., > 10000). In such cases, a threshold of > 1500 UMIs was enforced, and “true cells” were assigned again. Percentages of ribosomal, mitochondrial and *Malat1* counts were determined using the scanpy `calculate_qc_metrics()` function with “*Rps*”, “*Rpl*”, or “*mt*-” or “*Malat*” flags.

We performed QC with scanpy^130^ using median ± X*median absolute deviation filters (X = 4-5 for different metrics). Cut-offs were different for single-cell vs single-nucleus assays (single-nucleus also encompassing multiome samples). Specifically, these included:

percentage of mitochondrial counts (max_sc = 25, max_sn = 5, X_sc=5),
percentage of ribosomal counts (max_sc = 30, max_sn = 10, X_sc=5),
percentage of MALAT1 counts (max_sc = 30, X_sc=5),
percentage of counts in top 20 genes (X_sc=5, X_sn=5),
total counts (min_sc = 1000, min_sn = 1000),
genes detected (min_sc = 800, min_sn = 800),
log1p total counts (X_sc=4, X_sn=4),
log1p genes detected (X_sc=4, X_sn=4).

Filtering functions are implemented in the atlas module of the epitome_tools Python package to enable future atlasing efforts.

Following filtering, doublets were identified and removed using the Scrublet algorithm implemented in scanpy with default settings.

Ambient RNA was then removed using scAR^131^ for the top 200 most contaminating transcripts, as determined from cell-free droplets. We found scAR to overestimate the contribution of ambient transcripts (often entirely removing them during correction) and therefore applied shrinking (towards the median value of proportional contributions) to balance this effect. scAR was run for 600 epochs with a target of achieving 0.5% increased sparsity in the expression matrices. This left us with the original values for all transcripts, except for changes in the top 200 contaminating transcripts, often with most hormone transcripts in the top 10.

Following this, we performed an initial round of cell typing using CellAssign^132^ with curated cell type marker matrix of established pituitary markers. These were all taken from domain knowledge of pituitary physiology, main dot plots in publications, or from supplementary tables of highly significant marker genes in previous studies. Only genes that occurred at least twice across resources were used. Specifically, we used the following genes (in no specific order):

Stem_cells: *Sox2, Rbpms, Mia, Aqp3, Krt8, Krt18, Lcn2, Cyp2f2, Aldh1a2, Folr1, Pla2g7, Aldoc, Mgst1, Glul*
Corticotrophs: *Pomc, Crhr1, Tbx19, Gpc5, Tnt1, AW551984, Atp1a2*
Melanotrophs: *Pomc, Tbx19, Pax7, Oacyl, Pcsk2, Pkib, Megf11, Esm1, Etv1, Ascl1, Sparcl1*
Gonadotrophs: *Nr5a1, ENSMUSG00000100916.4, Fshb, Spp1, Tgfbr3l, Gnrhr*
Somatotrophs: *Pappa2, Car10, Rxrg, Gh, Ghrhr*
Lactotrophs: *Prl, Hepacam2, Edil3, Angpt1, Olfm1, Six6*
Thyrotroph: *Shox2, Dio2, Ttr, Rbp4, Trhr, Tshb*
Endothelial_cells: *Pecam1, Plvap, Igfbp7, Igfbp3, Emcn, Flt1*
Mesenchymal_cells: *Col1a1, Pdgfra, Ogn, Dcn, Inmt, Lum*
Pituicytes: *Fgf10, Rax, Scn7a, Gpc3, Nkx2-1*
Immune_cells: *Cd4, Tyrobp, Cd14, Trbc2, Cd68, C1qa, C1qc, Lyz2*
Erythrocytes: *Hba-a1, Hba-a2, Hbb-, Hba-x, Hbb-bs, Hbb-bt*

Following this coarse cell typing, all datasets were integrated using scVI. Initial cell type annotations were then smoothed using the 100 nearest neighbors of each cell obtained in the scVI latent space. Specifically, cell type values were adjusted if at least two-thirds of neighbors belonged to another cell type. This was repeated three times, reassigning the cell type of 52415 cells (about 5% of the RNA portion of the atlas). This step was applied to correct for dataset-specific inaccuracies introduced by CellAssign. Lastly, we noted two clusters in the scVI embedding which represented a mixture of cell types, indicative of low-quality cells that were retained past QC. We then found that most of these cells either come from the diphtheria toxin ablated samples^15,16,37^ marking truly dying cells or from IGVF samples, suggesting low quality cells using the PARSE technology. Cells from these clusters were removed from pseudobulking, and downstream analysis. However, these cells are still retained (to be as consistent with previous work as possible) in the individual datasets shared on *epitome*.

#### Sample inclusion criteria for analyses

Only samples with ≥300 cells were taken forward for any analysis. In addition, the 3 embryonic and 4 organoid samples were not used in any differential expression analysis. However all samples are available on the epitome platform.

From the IGVF samples three were filtered out as no pituitary cells were detected.

#### Accessing and processing chromatin accessibility datasets

Chromatin-accessibility data were obtained from Cell Ranger output files (specifically fragments.tsv files) from GEO. For one study,^19^ two samples were found to have been merged into a single count table. For these samples specifically, we realigned the reads using Cell Ranger ATAC 2.0. In another study, two donors had two biological replicates sequenced, but fragment files were only available for one in both cases.^35^ We accessed all four (2×2 replicates) and realigned these using Cell Ranger. All Cell Ranger fragment files were then loaded into SnapATAC2 ^133^ with the initial filter min_num_fragments > 1500.

We performed QC using a median ± 3*median absolute deviation filter, for log_n_frags. In addition, a hard cut-off of >7 was used for per cell transcription start site enrichment (TSSE). For each dataset, fragments were initially counted across genomic bins of 5000bp, and following the selection of the top 30000 features, Scrublet was used to identify and filter out doublets, leaving us high-quality cells.

Once high-quality cells were identified, peak calling was repeated with MACS3 ^134^ as implemented in SnapATAC2. Following this, peaks were filtered against the problematic regions from the ENCODE blacklist described in ^135^ using the file from https://github.com/Boyle-Lab/Blacklist/blob/master/lists/mm10-blacklist.v2.bed.gz with the SnapATAC2 function `snapatac2.pp.select_features()`. Notably the blacklist filtering only removed 3053 (about 1.14% of all) peaks, suggesting that blacklist regions are generally not problematic in snATAC-seq assays. The resulting peaks were then merged using a uniform 500 bp width, giving a Consensus Chromatin Landscape. Quantification was then performed by counting fragments (rather than reads, for reasons discussed in ^136^) falling in each peak. Peaks were retained if they were independently called by MACS3 in ≥10 datasets. Peaks in the Consensus

Chromatin Landscape were compared to dataset-specific peaks by counting the fragments in each set of peaks, and comparing them to fragments found in all genomic tiles/bins before peak calling (total fragment count).

For cell typing, we integrated all cells using poissonVI,^136^ and used cells from multiome studies as a bridge to transfer annotations previously made on the RNA level. Specifically, each cell was annotated by a “majority vote” system based on their 100 nearest neighbors in the poissonVI latent space. This step was repeated twice, such that all cells were then assigned to a given cell type. For robust analysis, we then pseudobulked each cell type in every sample using DecoupleR, such that each cell type was only pseudobulked if it had ≥100 cells in a given sample.

#### scVI and poissonVI models

To jointly analyze single cells from all transcriptomic datasets, we used the scVI variational auto-encoder framework.^137^ Specifically, we set each SRA_ID (one per dataset) as the batch_key, and set Comp_sex (computationally corrected sex assignment) and technologies (single-cell, single-nucleus, or multiome RNA) as categorical covariates. To increase mixing between technologies, percentages of mitochondrial, ribosomal, and intronic reads (using *Malat1* as proxy ^138^) were added as continuous covariates. Furthermore, to correct for age-dependent gene expression changes, we added “Age_numeric” as an additional continuous covariate. The model was set up using 2 hidden layers and a latent space with 30 dimensions. The generative end of the model was specified to be negative binomial, following considerations of the classic two-telegraph bursty transcription model. The model was then trained for 100 epochs (until convergence).

For chromatin accessibility datasets, we used the poissonVI variational auto-encoder framework,^136^ as implemented in scVI-tools.^137^ Here, we also set SRA_ID (one per dataset) as the batch_key, and set Sex and technologies (single-nucleus, or multiome ATAC) as categorical covariates. The model was set up using 2 hidden layers and a latent space with 30 dimensions, and a Poisson likelihood function (hence its name).

#### Dot plot construction

Generating dot plots for an entire atlas has the risk of introducing biases from samples with higher cell counts. To ensure that samples contribute evenly, we calculated the proportion of expression (dot plot circle size) and expression magnitude (dot plot circle color - calculated as mean log1p(CP10K)) per sample. We then calculated the mean proportion of expression across samples, and the mean of magnitude values. Because the mean values were calculated on a per-dataset basis, the aggregated means should not be biased by cell type abundance of individual datasets. These values were used to construct dot plots on the *epitome* platform, and all dot plots in the manuscript were exported from there, meaning they can also be reproduced by the reader.

#### Pseudobulk differential expression analysis

From each sc/snRNA-seq dataset, pseudobulk (summing all counts per cell type per dataset) versions were generated using decoupleR,^139^ which were merged in a final object. Only cell types with ≥50 cells were pseudobulked in each sample. We performed differential expression analysis for cell type markers and sex and age-dependent effects using limma voom.^140^ Similar workflows have been adopted by others using limma^141^ or pyDEseq2.^41^ Limma was chosen because it accommodates a mixed-effects model and also has a significant advantage in handling large datasets (>1000 pseudobulk samples in this case). Here each sample was normalized by its constituent number of cells, and then multiplied by the median number of cells across all samples. We used the `filterByExpr` function to remove low-expression genes. Then, TMM normalization within limma was used. Following this, a blocked design was pursued where different assay modalities (sc, sn, multiome) represented a block. The following model was used to retrieve marker genes: ∼ 0 + assignments.

In the analysis presented here, we defined “cell typing markers” as genes that might have high utility for annotating pituitary datasets in the future, or for planning spatial transcriptomics experiments. These were determined based on two criteria:

- Significantly higher expression in cell type X, compared to all other cell types (including non-anterior-pituitary cells, such as endothelial cells, pituicytes)
- Expression difference with a fold-change value of 4.

We also defined “lineage markers” which were evaluated only for cell types that belong in the anterior pituitary lineage, and comparisons were made for cell types at various branchpoints. These included 8 such comparisons, between:

Stem cells vs. Differentiated cells
Gonadotrophs vs. Stem cells and other differentiated cells
Corticotrophs/Melanotrophs vs Stem cells and other differentiated cells
Corticotrophs vs. Melanotrophs
Lactotrophs/Somatotrophs/Thyrotrophs vs Stem cells and other differentiated cells
Lactotrophs vs. Somatotrophs/Thyrotrophs
Somatotrophs vs. Lactotrophs/Thyrotrophs
Thytrotrophs vs. Somatotrophs/Lactotrophs

In each case, for a gene to be a lineage marker, it had to be significantly (FDR adjusted p-values < 0.05, and absolute log_2_FC > 0.5) different between the left-hand cell types compared to every single right-hand cell type separately.

For the number of samples used in each comparison, refer to “Quantification and statistical analysis”.

#### Correcting sex information

Following processing of the data, we checked whether the available meta-data on sex was consistent throughout transcriptomic datasets. To this end, we constructed a logistic regression model using scikit-learn^142^ on 3 sex-specific genes, the X-linked gene *Xist*, and two Y-linked genes *Ddx3y* and *Kdmd5*. Re-running the model on all samples highlighted outlier samples, that indicated switching of sexes. Two of these samples came from a publication that directly studied sex-specific effects.^18^

For chromatin accessibility datasets, we plotted fragment counts mapping to the Y chromosome and observed that sexes reported by authors matched the expected pattern, such that only male samples had fragments mapping to the Y chromosome. For these data, no corrections were required.

#### Age/Sex-dependent expression analysis

To identify age or sex-dependent gene expression changes the following models were used:

∼ 0 + assignments + assignments:Age_numeric and ∼ 0 + assignments + assignments:Sex. In the age-dependent analysis only single-cell (not nucleus/multiome) data was used. In the sex-dependent analysis a blocked design was applied to treat experimental methods as random effects. Hits with adjusted p-values < 0.05 (following Benjamini-Hochberg correction) were identified as statistically significant.

#### Gene expression box plot, and age-dependence scatter plot construction

Counts per million values for each transcript were calculated following TMM-correction in limma voom and normalizing counts to a million. Subsequently, a pseudocount of 1 was added to the count matrix and log_10_ transformation was performed. These values were used to construct box plots, and age-dependence scatter plots on the *epitome* platform. All such plots in the manuscript were exported from the epitome website, meaning they can also be reproduced by the reader. Specifically, the sex-dependence plots can be reproduced with the “Reproduce sex-specific analysis” option ticked, while the “Reproduce age-dependent analysis” setting allows reproducing the age-dependence plots.

#### Differential accessibility analysis

Pseudobulk analysis was then carried out in limma voom to identify cell type-specific open chromatin peaks. Limma voom was chosen here for the same reason as for the RNA-seq analysis. Each sample was normalized by its constituent number of cells, and then multiplied by the median number of cells across all samples. We used the `filterByExpr()` function to remove low-accessibility peaks. Then TMMwsp normalization within limma was used. The following model was used to retrieve marker peaks: ∼ 0 + assignments.

Lineage marking peaks were derived using the same methodology used for lineage marker genes.

Given the lack of samples with young or old ages, we did not test for accessibility changes with age, however, we were able to look at sex differences. This was performed with the following model ∼ 0 + assignments + assignments:Sex.

#### Gradient boosting models with XGBoost (Doublet Model and Cell Type Model)

The Doublet and Cell Type models were built using the XGBoost gradient boosting framework^57^, because of its ease of use and robustness. Before training, we normalized all datasets as log1p(CP10K).

For the Doublet Model, we performed feature selection by running XGBoost on one sample from every single study (except those held-out), following which we have identified the most important 1500 features across these results, using “`model.get_booster().get_score(importance_type=’weight’)`”. We then subsetted for these features as well as the previously identified Cell Typing markers.

Simulated doublets were generated for all samples separately, to avoid introducing doublets from pairs of cells coming from different experiments. Specifically, we looked for all cell types with ≥25 cells in a given sample, and sampled 150 pairs of cells with replacement for each pair of cells types. Importantly we did not simulate doublets coming from the same cell type (known as homotypic doublets), as these cannot be discriminated from true cells. Using this approach, we generated a balanced training dataset of 883,800 simulated doublets and 806,804 real cells.

We ran XGBoost with the following hyperparameters to fit models

~~~
params = {
   ’n_estimators’: 300,
   ’max_depth’: 12,
   ’subsample’: 0.8,
   “min_child_weight”: 2,
   ’learning_rate’:0.1,
   ’colsample_bytree’: 0.7,
   ’colsample_bylevel’: 0.7,
   ’colsample_bynode’: 0.7,
   ’objective’: ‘binary:logistic’,
   ’random_state’: 42,
   ’eval_metric’: ‘logloss’,
   ’tree_method’: “hist”
}
~~~

In addition, we used the `GroupKFold()` function in scikit-learn for cross-validation with 5 splits, and groups being source publications. We found this to be a more faithful representation of performance over splitting randomly on cells or even datasets, as there is some correlation across cells assayed in the same dataset and even in the same publication.

For later use, we took the model from the first-fold of the 5-fold cross validation. This is because we performed TPR/FPR calibration on this fold of the model, and the prediction threshold would have likely changed upon retraining the model on the entire atlas.

During cross-validation, we took the direct model output; however, for later evaluations in the manuscript we applied smoothing to cell type labels based on the closest 10 neighbors in PCA space. If ≥6 of the neighbors had the same initial cell type assignment, then that was assigned to the cell as a final label. This correction increased consistency on dimensionality reduction plots. In the evaluations we automatically set every missing gene (not present in the dataset as it was filtered out) to zero.

For evaluation, we picked three high quality whole-pituitary datasets from three different labs,^9,13,15^ and generated doublets comprising 10% of the total dataset. A 10% doublet rate is typical for single-cell experiments, we therefore created a fair setup for comparison with Scrublet. These doublets were simulated by randomly pairing cells of different cell types. Because of the random sampling we assumed that the resulting proportion of doublet types will reflect those in real experiments. In addition, this sampling approach allowed us to repeat this *in silico* experiment 20 times, to derive accurate estimates for the performance of the two approaches. Importantly, in each iteration we subsampled the dataset by 80%, and only then generated the 10% doublets. The 80% subsampling was introduced to sufficiently differentiate between rounds of simulations, particularly by introducing a large variability in rare cell types. We ran the scanpy implementation of Scrublet with default parameters on raw counts.

For the Cell Type Model, we performed feature selection by running XGBoost on one sample from every single study (except those held-out), following which we have identified the most important 1000 (3000 for ATAC) features across these results, using “`model.get_booster().get_score(importance_type=’weight’)`”. We then subsetted for these features as well as the previously identified Cell Typing markers. Importantly we removed hormone transcripts as these are typically the noisiest in every dataset.

We ran XGBoost with the following hyperparameters to fit models

~~~
params = {
   ’n_estimators’: 200,
   ’max_depth’: 8,
   ’subsample’: 0.8,
   “min_child_weight”:5,
   ’learning_rate’:0.05,
   ’colsample_bytree’: 0.7,
   ’colsample_bylevel’: 0.7,
   ’colsample_bynode’: 0.7,
   ’objective’: ‘multi:softmax’,
   ’num_class’: len(label_encoder.classes_),
   ’random_state’: 42,
   ’eval_metric’: ‘mlogloss’,
   ’tree_method’: “hist”
}
~~~

Here too, we used the `GroupKFold` function in scikit-learn for cross-validation with 5 splits, for reasons discussed above. The confusion matrix was calculated in the first fold of the cross-validation, however, for later evaluations we use a version of the model that was re-trained on the whole atlas (excluding held-out samples).

#### Doublet Model and Cell Type Model implementation

The models are implemented in the `workflow` module of epitome_tools Python package, which also facilitates cell typing and doublet detection on the epitome platform. We recommend users perform the joint workflow using:

~~~
from epitome_tools.workflow import celltype_doublet_workflow
annotated_adata = celltype_doublet_workflow(
                                        adata,
                                        active_assay=data_type,
                                        modality=“rna”
                                    )
~~~

Here active_assay can take the value of “sc”, “sn” or “multi_rna” for single-cell, single-nucleus and multiome data respectively, while modality can take the values “rna” and “atac”. Throughout all evaluations, we used this implementation of the model.

#### Unbiased manual annotation of cell types

To benchmark the Cell Type Model we compared its performance against manual annotation. Cell type annotation is typically done through iterations of unbiased clustering, plotting of known marker genes, and merging clusters, to annotate each cell type.

We tasked a co-author (O.S.) to analyze each dataset and return it with labels from the following categories: Corticotrophs, Gonadotrophs, Lactotrophs, Melanotrophs, Pituicytes, Somatotrophs, Stem cells, Thyrotrophs, Immune cells, Mesenchymal cells, Endothelial cells, Erythrocytes. We then ran the Cell Type Model with default settings and compared the predicted labels on the same cells.

#### Model evaluation and definitions

Throughout the manuscript we use various machine learning terms which are defined below.

True positive rate (TPR): The proportion of true cases that were identified as positive by the classifier.

False positive rate (TPR): The proportion of negative cases that were identified as positive by classifier.

Accuracy: Percentage of correct classifications.

Recall: Synonymous with the TPR, here used for multi-class classification. The proportion of true instances of a class that were correctly identified by the classifier.

Receiver operating characteristic (ROC) Curve: Varying the decision threshold of the classifier results in different pairs of TPR/FPR values. The ROC curve plots (TPR against FPR) all such pairs using each prediction probability as a potential threshold.

Area under the ROC curve: Metric used to evaluate classifiers, ranging between 0.5 and 1 for naive and perfect models respectively. The value can also be interpreted as the probability that the classifier will correctly distinguish between any pair of positive and negative cases. The roc curve was calculated using the `roc_curve()` function in scikit-learn.

Overlap in labels was calculated as the fraction of cells that share the same label over all cells. For the multiome data, this was evaluated only on cells that were present in both the ATAC and RNA parts of the data.

Confusion matrices were always row-normalized (e.g., divided by the total number of true values for a given class). Importantly, this returns recall values on the diagonal, which describe the number of cells that were correctly identified from each true class.

Model uncertainty was calculated as 1-P_max_, where P_max_ is the probability assigned to the most likely class. Using this definition, model uncertainty can range between 0 and 1-1/n_classes_ (in this case, 12 classes: ∼0.9167), as a fully uncertain model would predict all classes with the same probability.

#### Processing Bulk RNA-sequencing data - gene-level

Bulk RNA-sequencing datasets were retrieved as described for single-cell datasets. To be consistent with the single-cell results, the same reference genome (GRCm39), pseudoalignment software (kallisto-bustools) and differential expression analysis workflow (limma voom) were used. All bulk datasets are curated in **Supplementary Table 12**. Only those genes were visualized that reached significance in the CPA (except in Fig 3L, showing non-significant genes as well). Directional agreement was then evaluated for only those genes that reached significance in both the CPA and the bulk dataset as well. This ensured that only those genes are considered that had sufficient evidence on their direction in both datasets

#### Gene set enrichment

Enrichment of hit genes was performed using EnrichR with a set of *a priori* determined (to avoid cherry-picking results) databases: Reactome Pathways 2024, Wikipathways 2024 Mouse, and GO Biological Process 2023. All significantly enriched terms for stem cell lineage markers are found in **Supplementary Table 8**.

#### Ligand-receptor interaction prediction

Ligand-receptor interaction prediction was performed using LIANA+.^86^ This approach was chosen as it allows using an ensemble of ligand-receptor algorithms, as well as an integrated ligand-receptor interaction database. Here 6 algorithms (CellPhoneDB with 1000 permutations, CellChat with 1000 permutations, Connectome, log2FC, NATMI, SingleCellSignalR) were used as implemented in the `aggregate_rank()` function using a merged version of three databases (LIANA CellPhoneDB, LIANA CellChatDB, and CellPhoneDB 5.0.0). The difference between LIANA CellPhoneDB and CellPhoneDB 5.0.0 was that the latter was directly accessed from the CellPhoneDB website (https://www.cellphonedb.org/), however, both resources contained important interactions not covered by the other. By default LIANA+ performs robust rank aggregation across algorithms; however, it has no option to handle multiple datasets. We therefore run LIANA+ separately (so as not to bias results by dataset size) for all wild-type, whole-pituitary transcriptome datasets (n=156). This resulted in 156 lists of magnitude (how strong the expression is) and specificity (how unique this ligand-receptor signaling is across cell type pairs) values for all interactions. Following the LIANA+ step, individual lists of interactions were aggregated using a reimplementation of robust rank aggregation (RRA), which follows that in LIANA+, as well as in the original RRA paper ^87^. We used only whole-pituitary samples so as not to bias the results of robust rank aggregations with rankings from FACS-sorted datasets. Hits were called for genes with adjusted rho values (these are the outputs of RRA) for magnitude and specificity ranks < 0.05.

#### Annotating results as TFs and ligands

TF annotations were retrieved from the list of genes found at: https://esbl.nhlbi.nih.gov/Databases/KSBP2/Targets/Lists/TranscriptionFactors/. Ligands and receptors were annotated using the same curation that was used for the LIANA+ analysis.

#### TF motif enrichment

We performed enrichment for TF-binding motifs in differentially accessible peaks with Signac’s FindMotifs() function and using the JASPAR2022 database of TF binding motifs.

#### TF co-occurrence analysis

TF motif positions have been identified using the `AddMotifs()` Signac function using the “BSgenome.Mmusculus.UCSC.mm10” genome, and the JASPAR2022 CORE vertebrates motif collection. Following this, we have counted the co-occurrence of each motif alone and estimated the frequency of co-occurrence by chance. This was then compared to the observed co-occurrence using a Fisher’s exact test. As an example, for TFs “A” and “B”, we used a 2×2 contingency table with columns “A occurring” and “A not occurring”, and rows “B occurring” and “B not occurring”.

#### Mapping motifs to annotated genomic regions

Following extraction of motif positions, we used the annotatr package to assign genomic annotations to motif positions using the build_annotations() (with mm10 genome and the following annotation groups: ‘mm10_basicgenes’, ‘mm10_cpgs’, ‘mm10_genes_intergenic’, ‘mm10_genes_cds’, ‘mm10_genes_firstexons’, ‘mm10_genes_intronexonboundaries’, ‘mm10_genes_exonintronboundaries’, ‘mm10_lncrna_gencode’, ‘mm10_enhancers_fantom’) and annotate_regions() functions and summarized with the summarize_annotations() function.

#### RNAscope mRNA *in situ* hybridization

RNAscope mRNA *in situ* hybridization experiments were performed using the RNAscope 2.5 HD Duplex Reagent kit (ACD Bio #322430) as per manufacturer’s instructions. Stained sections were mounted in VectaMount Permanent Mounting Medium (Vector Laboratories, H-5000-60). The following probes were used: *Nrg1 (#418181), Erbb4 (#318721-C2), Slit2 (#449691), Robo1 (#475951-C2), Nr5a1 (#445731-C2), Nhlh2 (#527811), Sox2 (#401041-C2), Rfx4 (#520551), Six2 (#500011), Runx1 (#406671)*.

#### Immunofluorescence microscopy

Microscopy slides with mounted sections were deparaffinised and rehydrated through a descending graded ethanol series. Antigen retrieval was carried out using DeClere citrate retrieval buffer (pH 6.0) in a decloaking chamber NXGEN (Meraini Diagnostics) for 3 minutes at 110°C. Hydrophobic pap pen was applied around the sample and all incubation steps were carried out in a humified chamber.

Sections were incubated in blocking buffer (0.15% glycine, 2 mg/ml bovine serum albumine, 0.1% Triton-X in PBS), containing 10% serum (sheep or donkey) for 1 hr at room temperature. Sections were then incubated overnight with the respective primary antibodies at 4°C, in blocking buffer with 1% serum. Primary antibodies used were against SOX2 (1:300, R&D AF2018), LEF1 (1:100, Cell Signaling 2230S) and GAL (1:300, Invitrogen PA5-62069). Slides were washed in PBS with Triton-X 0.1% (PBST) then incubated with secondary antibodies in blocking buffer for 1 hr at room temperature (biotinylated anti-rabbit (1:350, Abcam ab207999), anti-goat Alexa fluor 488 (1:500, Abcam ab150129), anti-rabbit Alexa fluor 594 (1:500, Abcam ab150080)). Where appropriate, slides were washed using PBST and incubated with fluorophore-conjugated Streptavidin (1:500, Life Technologies S32355) for 1 hr at room temperature. Slides were then washed with PBST and incubated with Hoechst (1:10000, Life Technologies H3570). Slides were washed in PBST and mounted with VectaMount (Vector Laboratories, H1000).

#### Imaging

RNAscope slides were scanned with Nanozoomer-XR Digital slide scanner (Hamamatsu) and the images were processed using the Nanozoomer Digital Pathology (NDP) View software. In addition, some close-up images were taken with an Olympus BX34F Brightfield microscope using a 100X objective. Fluorescent stainings were imaged with a Leica TCS SP5 or a Zeiss LSM980 confocal microscope and the resulting images were processed using the Fiji software ^143^.

#### Quantification and statistical analysis

Statistical analyses were performed with dedicated software for bulk and single-cell genomics in either R or Python, as discussed above. For all analyses, we used a P-value (adjusted for multiple testing where appropriate) threshold of 0.05 to determine statistical significance. Multiple-testing corrections were applied as follows: Benjamini-Hochberg correction for marker discovery and sex-analysis, and Bonferroni corrections for age-analysis. In addition, we thresholded for absolute log_2_ fold-change in marker discovery (>0.5), age-analysis (>0.5 per log_10_ unit time) and sex-analysis (>1).

Throughout the text, we use the words biological replicates and samples interchangeably, and these refer to single-cell/nucleus samples. While some samples are whole pituitaries, some are sorted for a certain cell type. In each analysis, keeping in mind the statistical formulation of the question, these samples all function as replicates. We also count multiome samples as two replicates during our census presented in Figure 1 (1 RNA and 1 ATAC), as they contribute to both RNA and ATAC analyses, and are not treated as a single sample.

Sample sizes for the various differential expression/accessibility analyses were as follows (note that certain cell types might not have been present in sufficient numbers in every single biological replicate - e.g., 100 biological replicates might not mean 100 pseudobulks for cell type X):

CPA analyses (RNA) - do not include organoid and developmental samples:

Cell typing and lineage marker genes (all technologies, postnatal samples only): 210 samples and 18669 genes after filtering

Age-dependent expression (single-cell, not single-nucleus/multiome, postnatal samples only): 105 samples and 20595 genes after filtering

Sex-biased expression (10X, not PARSE, samples aged between P10-P150): 64 male vs 47 female samples and 18780 genes after filtering

CPA analyses (ATAC):

Lineage marker peaks (all samples): 39 samples and 226043 peaks after filtering

Sex-dependent peaks (P4 male multiome not used - only the rest of the samples that reached sexual maturity): 17 male vs 21 female samples and 225524 peaks after filtering

Bulk analyses:

Age-dependent genes: 4 young vs 4 aged samples and 15729 genes after filtering *hpg* mice sex-biased genes: 6 male vs 5 female samples and 15901 genes after filtering

Corticotroph sex-biased genes: 3 male vs 3 female samples and 16787 genes after filtering

Gonadotrophs sex-biased genes: 4 male vs. 4 female samples and 16594 genes after filtering

Whole-pituitary sex-biased genes: 26 male vs 29 female samples and 15510 genes after filtering

#### Additional resources

We have generated the electronic pituitary omics (*epitome*) platform, which is accessible at epitome-atlas.com.

## References

1. Lodge, E.J., Santambrogio, A., Russell, J.P., Xekouki, P., Jacques, T.S., Johnson, R.L., Thavaraj, S., Bornstein, S.R., and Andoniadou, C.L. (2019). Homeostatic and tumourigenic activity of SOX2+ pituitary stem cells is controlled by the LATS/YAP/TAZ cascade. eLife 8, e43996. 10.7554/eLife.43996.

2. Fauquier, T., Rizzoti, K., Dattani, M., Lovell-Badge, R., and Robinson, I.C.A.F. (2008). SOX2-expressing progenitor cells generate all of the major cell types in the adult mouse pituitary gland. Proc. Natl. Acad. Sci. U. S. A. 105, 2907–2912. 10.1073/pnas.0707886105.

3. Kelberman, D., Rizzoti, K., Avilion, A., Bitner-Glindzicz, M., Cianfarani, S., Collins, J., Chong, W.K., Kirk, J.M.W., Achermann, J.C., Ross, R., et al. (2006). Mutations within Sox2/SOX2 are associated with abnormalities in the hypothalamo-pituitary-gonadal axis in mice and humans. J. Clin. Invest. 116, 2442–2455. 10.1172/JCI28658.

4. Andoniadou, C.L., Matsushima, D., Mousavy Gharavy, S.N., Signore, M., Mackintosh, A.I., Schaeffer, M., Gaston-Massuet, C., Mollard, P., Jacques, T.S., Le Tissier, P., et al. (2013). Sox2+ Stem/Progenitor Cells in the Adult Mouse Pituitary Support Organ Homeostasis and Have Tumor-Inducing Potential. Cell Stem Cell 13, 433–445. 10.1016/j.stem.2013.07.004.

5. Li, S., Crenshaw, E.B., Rawson, E.J., Simmons, D.M., Swanson, L.W., and Rosenfeld, M.G. (1990). Dwarf locus mutants lacking three pituitary cell types result from mutations in the POU-domain gene pit-1. Nature 347, 528–533. 10.1038/347528a0.

6. Pulichino, A.-M., Vallette-Kasic, S., Tsai, J.P.-Y., Couture, C., Gauthier, Y., and Drouin, J. (2003). Tpit determines alternate fates during pituitary cell differentiation. Genes Dev. 17, 738–747. 10.1101/gad.1065703.

7. Ingraham, H.A., Lala, D.S., Ikeda, Y., Luo, X., Shen, W.H., Nachtigal, M.W., Abbud, R., Nilson, J.H., and Parker, K.L. (1994). The nuclear receptor steroidogenic factor 1 acts at multiple levels of the reproductive axis. Genes Dev. 8, 2302–2312. 10.1101/gad.8.19.2302.

8. Cheung, L.Y.M., George, A.S., McGee, S.R., Daly, A.Z., Brinkmeier, M.L., Ellsworth, B.S., and Camper, S.A. (2018). Single-Cell RNA Sequencing Reveals Novel Markers of Male Pituitary Stem Cells and Hormone-Producing Cell Types. Endocrinology 159, 3910–3924. 10.1210/en.2018-00750.

9. Ruf-Zamojski, F., Zhang, Z., Zamojski, M., Smith, G.R., Mendelev, N., Liu, H., Nudelman, G., Moriwaki, M., Pincas, H., Castanon, R.G., et al. (2021). Single nucleus multi-omics regulatory landscape of the murine pituitary. Nat. Commun. 12, 2677. 10.1038/s41467-021-22859-w.

10. Mayran, A., Sochodolsky, K., Khetchoumian, K., Harris, J., Gauthier, Y., Bemmo, A., Balsalobre, A., and Drouin, J. (2019). Pioneer and nonpioneer factor cooperation drives lineage specific chromatin opening. Nat. Commun. 10, 3807. 10.1038/s41467-019-11791-9.

11. Chen, Q., Leshkowitz, D., Blechman, J., and Levkowitz, G. (2020). Single-Cell Molecular and Cellular Architecture of the Mouse Neurohypophysis. eNeuro 7, ENEURO.0345-19.2019. 10.1523/ENEURO.0345-19.2019.

12. Ho, Y., Hu, P., Peel, M.T., Chen, S., Camara, P.G., Epstein, D.J., Wu, H., and Liebhaber, S.A. (2020). Single-cell transcriptomic analysis of adult mouse pituitary reveals sexual dimorphism and physiologic demand-induced cellular plasticity. Protein Cell 11, 565–583. 10.1007/s13238-020-00705-x.

13. Lopez, J.P., Brivio, E., Santambrogio, A., De Donno, C., Kos, A., Peters, M., Rost, N., Czamara, D., Brückl, T.M., Roeh, S., et al. (2021). Single-cell molecular profiling of all three components of the HPA axis reveals adrenal ABCB1 as a regulator of stress adaptation. Sci. Adv. 7, eabe4497. 10.1126/sciadv.abe4497.

14. Ruggiero-Ruff, R.E., Le, B.H., Villa, P.A., Lainez, N.M., Athul, S.W., Das, P., Ellsworth, B.S., and Coss, D. (2024). Single-Cell Transcriptomics Identifies Pituitary Gland Changes in Diet-Induced Obesity in Male Mice. Endocrinology 165, bqad196. 10.1210/endocr/bqad196.

15. Vennekens, A., Laporte, E., Hermans, F., Cox, B., Modave, E., Janiszewski, A., Nys, C., Kobayashi, H., Malengier-Devlies, B., Chappell, J., et al. (2021). Interleukin-6 is an activator of pituitary stem cells upon local damage, a competence quenched in the aging gland. Proc. Natl. Acad. Sci. U. S. A. 118, e2100052118. 10.1073/pnas.2100052118.

16. Laporte, E., Hermans, F., De Vriendt, S., Vennekens, A., Lambrechts, D., Nys, C., Cox, B., and Vankelecom, H. (2022). Decoding the activated stem cell phenotype of the neonatally maturing pituitary. eLife 11, e75742. 10.7554/eLife.75742.

17. Li, Y., Wang, J., Wang, R., Chang, Y., and Wang, X. (2023). Gut bacteria induce IgA expression in pituitary hormone-secreting cells during aging. iScience 26, 107747. 10.1016/j.isci.2023.107747.

18. Miles, T.K., Odle, A.K., Byrum, S.D., Lagasse, A., Haney, A., Ortega, V.G., Bolen, C.R., Banik, J., Reddick, M.M., Herdman, A., et al. (2023). Anterior Pituitary Transcriptomics Following a High-Fat Diet: Impact of Oxidative Stress on Cell Metabolism. Endocrinology 165, bqad191. 10.1210/endocr/bqad191.

19. Bohaczuk, S.C., Thackray, V.G., Shen, J., Skowronska-Krawczyk, D., and Mellon, P.L. (2021). FSHB Transcription is Regulated by a Novel 5′ Distal Enhancer With a Fertility-Associated Single Nucleotide Polymorphism. Endocrinology 162, bqaa181. 10.1210/endocr/bqaa181.

20. Schang, G., Ongaro, L., Brûlé, E., Zhou, X., Wang, Y., Boehm, U., Ruf-Zamojski, F., Zamojski, M., Mendelev, N., Seenarine, N., et al. (2022). Transcription factor GATA2 may potentiate follicle-stimulating hormone production in mice *via* induction of the BMP antagonist gremlin in gonadotrope cells. J. Biol. Chem. 298, 102072. 10.1016/j.jbc.2022.102072.

21. Lin, Y.-F., Schang, G., Buddle, E.R.S., Schultz, H., Willis, T.L., Ruf-Zamojski, F., Zamojski, M., Mendelev, N., Boehm, U., Sealfon, S.C., et al. (2022). Steroidogenic Factor 1 Regulates Transcription of the Inhibin B Coreceptor in Pituitary Gonadotrope Cells. Endocrinology 163, bqac131. 10.1210/endocr/bqac131.

22. Rizzoti, K., Chakravarty, P., Sheridan, D., and Lovell-Badge, R. SOX9-positive pituitary stem cells differ according to their position in the gland and maintenance of their progeny depends on context. Sci. Adv. 9, eadf6911. 10.1126/sciadv.adf6911.

23. Cheung, L.Y.M., Menage, L., Rizzoti, K., Hamilton, G., Dumontet, T., Basham, K., Daly, A.Z., Brinkmeier, M.L., Masser, B.E., Treier, M., et al. (2023). Novel Candidate Regulators and Developmental Trajectory of Pituitary Thyrotropes. Endocrinology 164, bqad076. 10.1210/endocr/bqad076.

24. Allensworth-James, M., Banik, J., Odle, A., Hardy, L., Lagasse, A., Moreira, A.R.S., Bird, J., Thomas, C.L., Avaritt, N., Kharas, M.G., et al. (2021). Control of the Anterior Pituitary Cell Lineage Regulator POU1F1 by the Stem Cell Determinant Musashi. Endocrinology 162, bqaa245. 10.1210/endocr/bqaa245.

25. Moncho-Amor, V., Chakravarty, P., Galichet, C., Matheu, A., Lovell-Badge, R., and Rizzoti, K. (2021). SOX2 is required independently in both stem and differentiated cells for pituitary tumorigenesis in p27-null mice. Proc. Natl. Acad. Sci. U. S. A. 118, e2017115118. 10.1073/pnas.2017115118.

26. Bastedo, W.E., Scott, R.W., Arostegui, M., and Underhill, T.M. (2024). Single-cell analysis of mesenchymal cells in permeable neural vasculature reveals novel diverse subpopulations of fibroblasts. Fluids Barriers CNS 21, 31. 10.1186/s12987-024-00535-7.

27. Zhang, Z., Ruf-Zamojski, F., Zamojski, M., Bernard, D.J., Chen, X., Troyanskaya, O.G., and Sealfon, S.C. (2024). Peak-agnostic high-resolution cis-regulatory circuitry mapping using single cell multiome data. Nucleic Acids Res. 52, 572–582. 10.1093/nar/gkad1166.

28. Cheung, L.Y.M., and Camper, S.A. (2020). PROP1-Dependent Retinoic Acid Signaling Regulates Developmental Pituitary Morphogenesis and Hormone Expression. Endocrinology 161, bqaa002. 10.1210/endocr/bqaa002.

29. Masser, B.E., Brinkmeier, M.L., Lin, Y., Liu, Q., Miyazaki, A., Nayeem, J., and Cheung, L.Y.M. (2024). Gene Misexpression in a Smoc2+ve/Sox2-Low Population in Juvenile Prop1-Mutant Pituitary Gland. J. Endocr. Soc. 8, bvae146. 10.1210/jendso/bvae146.

30. Qian, Q., Li, M., Zhang, Z., Davis, S.W., Rahmouni, K., Norris, A.W., Cao, H., Ding, W.-X., Hotamisligil, G.S., and Yang, L. (2024). Obesity disrupts the pituitary-hepatic UPR communication leading to NAFLD progression. Cell Metab. 36, 1550–1565.e9. 10.1016/j.cmet.2024.04.014.

31. Martinez-Mayer, J., Brinkmeier, M.L., O’Connell, S.P., Ukagwu, A., Marti, M.A., Miras, M., Forclaz, M.V., Benzrihen, M.G., Cheung, L.Y.M., Camper, S.A., et al. (2024). Knockout mice with pituitary malformations help identify human cases of hypopituitarism. Genome Med. 16, 75. 10.1186/s13073-024-01347-y.

32. Kang, Y., Lee, S., Kim, Y., Lee, E., Oh, C., and Ku, C. (2023). A single-cell transcriptomic atlas of mouse pituitary aging. (Gene Expression Omnibus). https://www.ncbi.nlm.nih.gov/geo/query/acc.cgi?acc=GSE242296 https://www.ncbi.nlm.nih.gov/geo/query/acc.cgi?acc=GSE242296.

33. Sheridan, D., Chakravarty, P., Golan, G., Shiakola, Y., Olsen, J., Burnett, E., Galichet, C., Fiordelisio, T., Mollard, P., Melamed, P., et al. (2025). Gonadotrophs have a dual origin, with most derived from early postnatal pituitary stem cells. Nat. Commun. 16, 4280. 10.1038/s41467-025-59495-7.

34. Khetchoumian, K., Sochodolsky, K., Lafont, C., Gouhier, A., Bemmo, A., Kherdjemil, Y., Kmita, M., Le Tissier, P., Mollard, P., Christian, H., et al. (2024). Paracrine FGF1 signaling directs pituitary architecture and size. Proc. Natl. Acad. Sci. 121, e2410269121. 10.1073/pnas.2410269121.

35. Wang, Y., Thistlethwaite, W., Tadych, A., Ruf-Zamojski, F., Bernard, D.J., Cappuccio, A., Zaslavsky, E., Chen, X., Sealfon, S.C., and Troyanskaya, O.G. (2024). Automated single-cell omics end-to-end framework with data-driven batch inference. Cell Syst. 15, 982–990.e5. 10.1016/j.cels.2024.09.003.

36. Huang, Y., Wang, Q., Zhou, W., Jiang, Y., He, K., Huang, W., Feng, Y., Wu, H., Liu, L., Pan, Y., et al. (2024). Prenatal p25-activated Cdk5 induces pituitary tumorigenesis through MCM2 phosphorylation-mediated cell proliferation. Neoplasia N. Y. N 57, 101054. 10.1016/j.neo.2024.101054.

37. Vriendt, S.D., Laporte, E., Abaylı, B., Hoekx, J., Hermans, F., Lambrechts, D., and Vankelecom, H. (2025). Single-cell transcriptome atlas of male mouse pituitary across postnatal life highlighting its stem cell landscape. iScience 28. 10.1016/j.isci.2024.111708.

38. Ongaro, L., Zhou, X., Wang, Y., Schultz, H., Zhou, Z., Buddle, E.R.S., Brûlé, E., Lin, Y.-F., Schang, G., Hagg, A., et al. (2025). Muscle-derived myostatin is a major endocrine driver of follicle-stimulating hormone synthesis. Science 387, 329–336. 10.1126/science.adi4736.

39. Brinkmeier, M.L., Cheung, L.Y.M., O’Connell, S.P., Gutierrez, D.K., Rhoads, E.C., Camper, S.A., and Davis, S.W. (2025). Nucleoredoxin regulates WNT signaling during pituitary stem cell differentiation. Hum. Mol. Genet., ddaf032. 10.1093/hmg/ddaf032.

40. Miles, T.K., Odle, A.K., Byrum, S.D., Lagasse, A.N., Haney, A.C., Ortega, V.G., Herdman, A.K., MacNicol, M.C., MacNicol, A.M., and Childs, G.V. (2025). Ablation of Leptin Receptor Signaling Alters Somatotrope Transcriptome Maturation in Female Mice. Endocrinology, bqaf036. 10.1210/endocr/bqaf036.

41. Rebboah, E., Weber, R., Abdollahzadeh, E., Swarna, N., Sullivan, D.K., Trout, D., Reese, F., Liang, H.Y., Filimban, G., Mahdipoor, P., et al. (2025). Systematic cell-type resolved transcriptomes of 8 tissues in 8 lab and wild-derived mouse strains captures global and local expression variation. bioRxiv, 2025.04.21.649844. 10.1101/2025.04.21.649844.

42. Cheung, L. (2025). Fundamental mechanisms causing pituitary stem cell aging in mice and humans. (Gene Expression Omnibus). https://www.ncbi.nlm.nih.gov/geo/query/acc.cgi?acc=GSE299835 https://www.ncbi.nlm.nih.gov/geo/query/acc.cgi?acc=GSE299835.

43. Wei, R., Du, Z., Tao, W., and Zhang, C. (2025). Single-cell transcriptomic analysis reveals that inflammation drives the unfolded protein response during endocrine aging in mice. (Gene Expression Omnibus). https://www.ncbi.nlm.nih.gov/geo/query/acc.cgi?acc=GSE239316 https://www.ncbi.nlm.nih.gov/geo/query/acc.cgi?acc=GSE239316.

44. Rich, J.M., Moses, L., Einarsson, P.H., Jackson, K., Luebbert, L., Booeshaghi, A.S., Antonsson, S., Sullivan, D.K., Bray, N., Melsted, P., et al. (2024). The impact of package selection and versioning on single-cell RNA-seq analysis. Preprint at bioRxiv, 10.1101/2024.04.04.588111 10.1101/2024.04.04.588111.

45. Wilks, C., Zheng, S.C., Chen, F.Y., Charles, R., Solomon, B., Ling, J.P., Imada, E.L., Zhang, D., Joseph, L., Leek, J.T., et al. (2021). recount3: summaries and queries for large-scale RNA-seq expression and splicing. Genome Biol. 22, 323. 10.1186/s13059-021-02533-6.

46. Lachmann, A., Torre, D., Keenan, A.B., Jagodnik, K.M., Lee, H.J., Wang, L., Silverstein, M.C., and Ma’ayan, A. (2018). Massive mining of publicly available RNA-seq data from human and mouse. Nat. Commun. 9, 1366. 10.1038/s41467-018-03751-6.

47. Youngblut, N.D., Carpenter, C., Prashar, J., Ricci-Tam, C., Ilango, R., Teyssier, N., Konermann, S., Hsu, P.D., Dobin, A., Burke, D.P., et al. (2025). scBaseCount: an AI agent-curated, uniformly processed, and continually expanding single cell data repository. Preprint at bioRxiv, 10.1101/2025.02.27.640494 10.1101/2025.02.27.640494.

48. Booeshaghi, A.S., Galvez-Merchán, Á., and Pachter, L. (2024). Algorithms for a Commons Cell Atlas. Preprint at bioRxiv, 10.1101/2024.03.23.586413 10.1101/2024.03.23.586413.

49. Kövér, B., Kaufman-Cook, J., Sherwin, O., Vazquez Segoviano, M., Kemkem, Y., Lu, H.-C., and Andoniadou, C.L. (2025). Electronic Pituitary Omics (epitome) platform. (Zenodo). 10.5281/zenodo.17154161 10.5281/zenodo.17154161.

50. Mukamel, E.A., and Yu, Z. (2025). False positives in study of memory-related gene expression. Nature 642, E1–E3. 10.1038/s41586-025-08988-y.

51. Murphy, A.E., and Skene, N.G. (2022). A balanced measure shows superior performance of pseudobulk methods in single-cell RNA-sequencing analysis. Nat. Commun. 13, 7851. 10.1038/s41467-022-35519-4.

52. Thurman, A.L., Ratcliff, J.A., Chimenti, M.S., and Pezzulo, A.A. (2021). Differential gene expression analysis for multi-subject single-cell RNA-sequencing studies with aggregateBioVar. Bioinformatics 37, 3243–3251. 10.1093/bioinformatics/btab337.

53. Squair, J.W., Gautier, M., Kathe, C., Anderson, M.A., James, N.D., Hutson, T.H., Hudelle, R., Qaiser, T., Matson, K.J.E., Barraud, Q., et al. (2021). Confronting false discoveries in single-cell differential expression. Nat. Commun. 12, 5692. 10.1038/s41467-021-25960-2.

54. Crowell, H.L., Soneson, C., Germain, P.-L., Calini, D., Collin, L., Raposo, C., Malhotra, D., and Robinson, M.D. (2020). muscat detects subpopulation-specific state transitions from multi-sample multi-condition single-cell transcriptomics data. Nat. Commun. 11, 6077. 10.1038/s41467-020-19894-4.

55. Junttila, S., Smolander, J., and Elo, L.L. (2022). Benchmarking methods for detecting differential states between conditions from multi-subject single-cell RNA-seq data. Brief. Bioinform. 23, bbac286. 10.1093/bib/bbac286.

56. Hafemeister, C., and Halbritter, F. (2023). Single-cell RNA-seq differential expression tests within a sample should use pseudo-bulk data of pseudo-replicates. Preprint at bioRxiv, 10.1101/2023.03.28.534443 10.1101/2023.03.28.534443.

57. Chen, T., and Guestrin, C. (2016). XGBoost: A Scalable Tree Boosting System. In Proceedings of the 22nd ACM SIGKDD International Conference on Knowledge Discovery and Data Mining, pp. 785–794. 10.1145/2939672.2939785.

58. Wolock, S.L., Lopez, R., and Klein, A.M. (2019). Scrublet: Computational Identification of Cell Doublets in Single-Cell Transcriptomic Data. Cell Syst. 8, 281–291.e9. 10.1016/j.cels.2018.11.005.

59. Hou, H., Chan, C., Yuki, K.E., Sokolowski, D., Roy, A., Qu, R., Uusküla-Reimand, L., Faykoo-Martinez, M., Hudson, M., Corre, C., et al. (2022). Postnatal developmental trajectory of sex-biased gene expression in the mouse pituitary gland. Biol. Sex Differ. 13, 57. 10.1186/s13293-022-00467-7.

60. Hsu, D.W., Hooi, S.C., Hedley-Whyte, E.T., Strauss, R.M., and Kaplan, L.M. (1991). Coexpression of galanin and adrenocorticotropic hormone in human pituitary and pituitary adenomas. Am. J. Pathol. 138, 897–909.

61. Leung, B., Iisma, T.P., Leung, K.-C., Hort, Y.J., Turner, J., Sheehy, J.P., and Ho, K.K.Y. (2002). Galanin in human pituitary adenomas: frequency and clinical significance. Clin. Endocrinol. (Oxf.) 56, 397–403. 10.1046/j.1365-2265.2002.01486.x.

62. Wynick, D., Small, C.J., Bacon, A., Holmes, F.E., Norman, M., Ormandy, C.J., Kilic, E., Kerr, N.C.H., Ghatei, M., Talamantes, F., et al. (1998). Galanin regulates prolactin release and lactotroph proliferation. Proc. Natl. Acad. Sci. 95, 12671–12676. 10.1073/pnas.95.21.12671.

63. Cai, A., Bowers, R.C., Moore, J.P., Jr., and Hyde, J.F. (1998). Function of Galanin in the Anterior Pituitary of Estrogen-Treated Fischer 344 Rats: Autocrine and Paracrine Regulation of Prolactin Secretion*. Endocrinology 139, 2452–2458. 10.1210/endo.139.5.6025.

64. Shima, Y., Miyabayashi, K., Mori, T., Ono, K., Kajimoto, M., Cho, H.L., Tsuchida, H., Uenoyama, Y., Tsukamura, H., Suzuki, K., et al. (2022). Intronic Enhancer Is Essential for Nr5a1 Expression in the Pituitary Gonadotrope and for Postnatal Development of Male Reproductive Organs in a Mouse Model. Int. J. Mol. Sci. 24, 192. 10.3390/ijms24010192.

65. Duncan, P.J., Romanò, N., Nair, S.V., Murray, J.F., Le Tissier, P., and Shipston, M.J. (2023). Sex differences in pituitary corticotroph excitability. Front. Physiol. 14. 10.3389/fphys.2023.1205162.

66. Iacovazzo, D., Caswell, R., Bunce, B., Jose, S., Yuan, B., Hernández-Ramírez, L.C., Kapur, S., Caimari, F., Evanson, J., Ferraù, F., et al. (2016). Germline or somatic GPR101 duplication leads to X-linked acrogigantism: a clinico-pathological and genetic study. Acta Neuropathol. Commun. 4, 56. 10.1186/s40478-016-0328-1.

67. Alonso, C.A.I., David, C.D., Toufaily, C., Wang, Y., Zhou, X., Ongaro, L., Nudelman, G., Nair, V.D., Ruf-Zamojski, F., Boehm, U., et al. (2023). Activating Transcription Factor 3 Stimulates Follicle-Stimulating Hormone-β Expression In Vitro But Is Dispensable for Follicle-Stimulating Hormone Production in Murine Gonadotropes In Vivo. Endocrinology 164, bqad050. 10.1210/endocr/bqad050.

68. Funk, M.C., Gleixner, J.G., Heigwer, F., Vonficht, D., Valentini, E., Aydin, Z., Tonin, E., Prete, S.D., Mahara, S., Throm, Y., et al. (2023). Aged intestinal stem cells propagate cell-intrinsic sources of inflammaging in mice. Dev. Cell 58, 2914–2929.e7. 10.1016/j.devcel.2023.11.013.

69. Gonzalez-Meljem, J.M., Haston, S., Carreno, G., Apps, J.R., Pozzi, S., Stache, C., Kaushal, G., Virasami, A., Panousopoulos, L., Mousavy-Gharavy, S.N., et al. (2017). Stem cell senescence drives age-attenuated induction of pituitary tumours in mouse models of paediatric craniopharyngioma. Nat. Commun. 8, 1819. 10.1038/s41467-017-01992-5.

70. Gonzalez-Meljem, J.M., Cao, L., Apps, J.R., and Martinez-Barbera, J.P. (2025). Decoding craniopharyngioma: From mechanisms to therapy. Best Pract. Res. Clin. Endocrinol. Metab. 39, 102051. 10.1016/j.beem.2025.102051.

71. Saul, D., Kosinsky, R.L., Atkinson, E.J., Doolittle, M.L., Zhang, X., LeBrasseur, N.K., Pignolo, R.J., Robbins, P.D., Niedernhofer, L.J., Ikeno, Y., et al. (2022). A new gene set identifies senescent cells and predicts senescence-associated pathways across tissues. Nat. Commun. 13, 4827. 10.1038/s41467-022-32552-1.

72. Lopes-Pinto, M., Lacerda-Nobre, E., Silva, A.L., and Marques, P. (2024). Therapeutical Usefulness of PD-1/PD-L1 Inhibitors in Aggressive or Metastatic Pituitary Tumours. Cancers 16, 3033. 10.3390/cancers16173033.

73. Cossu, G., La Rosa, S., Brouland, J.P., Pitteloud, N., Harel, E., Santoni, F., Brunner, M., Daniel, R.T., and Messerer, M. (2023). PD-L1 Expression in Pituitary Neuroendocrine Tumors/Pituitary Adenomas. Cancers 15, 4471. 10.3390/cancers15184471.

74. Leu, J.-S., Chang, S.-Y., Mu, C.-Y., Chen, M.-L., and Yan, B.-S. (2018). Functional domains of SP110 that modulate its transcriptional regulatory function and cellular translocation. J. Biomed. Sci. 25, 34. 10.1186/s12929-018-0434-4.

75. Fox, G.J., Sy, D.N., Nhung, N.V., Yu, B., Ellis, M.K., Van Hung, N., Cuong, N.K., Thi Lien, L., Marks, G.B., Saunders, B.M., et al. (2014). Polymorphisms of SP110 are associated with both pulmonary and extra-pulmonary tuberculosis among the Vietnamese. PloS One 9, e99496. 10.1371/journal.pone.0099496.

76. Cui, X., Yuan, T., Ning, P., Han, J., Liu, Y., Feng, J., Lian, F., Hao, M., Dong, L., Hao, J., et al. (2022). Polymorphisms in the ASAP1 and SP110 Genes and Its Association with the Susceptibility to Pulmonary Tuberculosis in a Mongolian Population. J. Immunol. Res. 2022, 2713869. 10.1155/2022/2713869.

77. Carey, K.L., Paulus, G.L.C., Wang, L., Balce, D.R., Luo, J.W., Bergman, P., Ferder, I.C., Kong, L., Renaud, N., Singh, S., et al. (2020). TFEB Transcriptional Responses Reveal Negative Feedback by BHLHE40 and BHLHE41. Cell Rep. 33, 108371. 10.1016/j.celrep.2020.108371.

78. Kreslavsky, T., Vilagos, B., Tagoh, H., Poliakova, D.K., Schwickert, T.A., Wöhner, M., Jaritz, M., Weiss, S., Taneja, R., Rossner, M.J., et al. (2017). Essential role for the transcription factor Bhlhe41 in regulating the development, self-renewal and BCR repertoire of B-1a cells. Nat. Immunol. 18, 442–455. 10.1038/ni.3694.

79. Wu, W., Cogan, J.D., Pfäffle, R.W., Dasen, J.S., Frisch, H., O’Connell, S.M., Flynn, S.E., Brown, M.R., Mullis, P.E., Parks, J.S., et al. (1998). Mutations in PROP1 cause familial combined pituitary hormone deficiency. Nat. Genet. 18, 147–149. 10.1038/ng0298-147.

80. Pérez Millán, M.I., Brinkmeier, M.L., Mortensen, A.H., and Camper, S.A. (2016). PROP1 triggers epithelial-mesenchymal transition-like process in pituitary stem cells. eLife 5, e14470. 10.7554/eLife.14470.

81. Russell, J.P., Lim, X., Santambrogio, A., Yianni, V., Kemkem, Y., Wang, B., Fish, M., Haston, S., Grabek, A., Hallang, S., et al. (2021). Pituitary stem cells produce paracrine WNT signals to control the expansion of their descendant progenitor cells. eLife 10, e59142. 10.7554/eLife.59142.

82. Zhu, X., Tollkuhn, J., Taylor, H., and Rosenfeld, M.G. (2015). Notch-Dependent Pituitary SOX2+ Stem Cells Exhibit a Timed Functional Extinction in Regulation of the Postnatal Gland. Stem Cell Rep. 5, 1196–1209. 10.1016/j.stemcr.2015.11.001.

83. Ge, X., Weis, K., and Raetzman, L. (2024). Glycoprotein hormone subunit alpha 2 (GPHA2): A pituitary stem cell-expressed gene associated with NOTCH2 signaling. Mol. Cell. Endocrinol. 586, 112163. 10.1016/j.mce.2024.112163.

84. Xekouki, P., Lodge, E.J., Matschke, J., Santambrogio, A., Apps, J.R., Sharif, A., Jacques, T.S., Aylwin, S., Prevot, V., Li, R., et al. (2019). Non-secreting pituitary tumours characterised by enhanced expression of YAP/TAZ. Endocr. Relat. Cancer 26, 215–225. 10.1530/ERC-18-0330.

85. Lodge, E.J., Russell, J.P., Patist, A.L., Francis-West, P., and Andoniadou, C.L. (2016). Expression Analysis of the Hippo Cascade Indicates a Role in Pituitary Stem Cell Development. Front. Physiol. 7, 114. 10.3389/fphys.2016.00114.

86. Dimitrov, D., Schäfer, P.S.L., Farr, E., Rodriguez-Mier, P., Lobentanzer, S., Badia-i-Mompel, P., Dugourd, A., Tanevski, J., Ramirez Flores, R.O., and Saez-Rodriguez, J. (2024). LIANA+ provides an all-in-one framework for cell–cell communication inference. Nat. Cell Biol. 26, 1613–1622. 10.1038/s41556-024-01469-w.

87. Kolde, R., Laur, S., Adler, P., and Vilo, J. (2012). Robust rank aggregation for gene list integration and meta-analysis. Bioinformatics 28, 573–580. 10.1093/bioinformatics/btr709.

88. Willis, T.L., Lodge, E.J., Andoniadou, C.L., and Yianni, V. (2022). Cellular interactions in the pituitary stem cell niche. Cell. Mol. Life Sci. 79, 612. 10.1007/s00018-022-04612-8.

89. Le Tissier, P.R., Murray, J.F., and Mollard, P. (2022). A New Perspective on Regulation of Pituitary Plasticity: The Network of SOX2-Positive Cells May Coordinate Responses to Challenge. Endocrinology 163, bqac089. 10.1210/endocr/bqac089.

90. Budry, L., Balsalobre, A., Gauthier, Y., Khetchoumian, K., L’Honoré, A., Vallette, S., Brue, T., Figarella-Branger, D., Meij, B., and Drouin, J. (2012). The selector gene Pax7 dictates alternate pituitary cell fates through its pioneer action on chromatin remodeling. Genes Dev. 26, 2299–2310. 10.1101/gad.200436.112.

91. Budry, L., Couture, C., Balsalobre, A., and Drouin, J. (2011). The Ets Factor Etv1 Interacts with Tpit Protein for Pituitary Pro-opiomelanocortin (POMC) Gene Transcription. J. Biol. Chem. 286, 25387–25396. 10.1074/jbc.M110.202788.

92. Lamolet, B., Poulin, G., Chu, K., Guillemot, F., Tsai, M.-J., and Drouin, J. (2004). Tpit-independent function of NeuroD1(BETA2) in pituitary corticotroph differentiation. Mol. Endocrinol. Baltim. Md 18, 995–1003. 10.1210/me.2003-0127.

93. Poulin, G., Lebel, M., Chamberland, M., Paradis, F.W., and Drouin, J. (2000). Specific Protein-Protein Interaction between Basic Helix-Loop-Helix Transcription Factors and Homeoproteins of the Pitx Family. Mol. Cell. Biol. 20, 4826–4837. 10.1128/mcb.20.13.4826-4837.2000.

94. Poulin, G., Turgeon, B., and Drouin, J. (1997). NeuroD1/beta2 contributes to cell-specific transcription of the proopiomelanocortin gene. Mol. Cell. Biol. 17, 6673–6682. 10.1128/MCB.17.11.6673.

95. Oyama, K., Sanno, N., Teramoto, A., and Osamura, R.Y. (2001). Expression of Neuro D1 in Human Normal Pituitaries and Pituitary Adenomas. Mod. Pathol. 14, 892–899. 10.1038/modpathol.3880408.

96. Yoshizawa, T., Handa, Y., Uematsu, Y., Takeda, S., Sekine, K., Yoshihara, Y., Kawakami, T., Arioka, K., Sato, H., Uchiyama, Y., et al. (1997). Mice lacking the vitamin D receptor exhibit impaired bone formation, uterine hypoplasia and growth retardation after weaning. Nat. Genet. 16, 391–396. 10.1038/ng0897-391.

97. Song, Y., Fleet, J.C., and Kato, S. (2003). Vitamin D Receptor (VDR) Knockout Mice Reveal VDR-Independent Regulation of Intestinal Calcium Absorption and ECaC2 and Calbindin D9k mRNA. J. Nutr. 133, 374–380. 10.1093/jn/133.2.374.

98. Sánchez-Pacheco, A., Palomino, T., and Aranda, A. (1995). Retinoic acid induces expression of the transcription factor GHF-1/Pit-1 in pituitary prolactin- and growth hormone-producing cell lines. Endocrinology 136, 5391–5398. 10.1210/endo.136.12.7588287.

99. Bedo, G., Santisteban, P., and Aranda, A. (1989). Retinoic acid regulates growth hormone gene expression. Nature 339, 231–234. 10.1038/339231a0.

100. Orlov, I., Rochel, N., Moras, D., and Klaholz, B.P. (2012). Structure of the full human RXR/VDR nuclear receptor heterodimer complex with its DR3 target DNA. EMBO J. 31, 291–300. 10.1038/emboj.2011.445.

101. Seoane, S., and Perez-Fernandez, R. (2006). The Vitamin D Receptor Represses Transcription of the Pituitary Transcription Factor Pit-1 Gene without Involvement of the Retinoid X Receptor. Mol. Endocrinol. 20, 735–748. 10.1210/me.2005-0253.

102. Yu, H., Guo, B., Miao, Z., Chen, C., Song, Y., and Yang, J. (2025). A high-fat diet suppresses growth hormone synthesis and secretion by influencing the Vit D receptor and Pit1. Endocrine 89, 508–520. 10.1007/s12020-025-04270-3.

103. Arnhold, I.J.P., França, M.M., Carvalho, L.R., Mendonca, B.B., and Jorge, A.A.L. (2015). Role of GLI2 in hypopituitarism phenotype. J. Mol. Endocrinol. 54, R141–150. 10.1530/JME-15-0009.

104. Babu, D., Fanelli, A., Mellone, S., Muniswamy, R., Wasniewska, M., Prodam, F., Petri, A., Bellone, S., Salerno, M.C., and Giordano, M. (2019). Novel GLI2 mutations identified in patients with Combined Pituitary Hormone Deficiency (CPHD): Evidence for a pathogenic effect by functional characterization. Clin. Endocrinol. (Oxf.) 90, 449–456. 10.1111/cen.13914.

105. Stallings, C.E., Das, P., Athul, S.W., Ukagwu, A.E., Jensik, P.J., and Ellsworth, B.S. (2024). FOXO1 regulates expression of *Neurod4* in the pituitary gland. Mol. Cell. Endocrinol. 583, 112128. 10.1016/j.mce.2023.112128.

106. Kapali, J., Kabat, B.E., Schmidt, K.L., Stallings, C.E., Tippy, M., Jung, D.O., Edwards, B.S., Nantie, L.B., Raeztman, L.T., Navratil, A.M., et al. (2016). Foxo1 Is Required for Normal Somatotrope Differentiation. Endocrinology 157, 4351–4363. 10.1210/en.2016-1372.

107. Das, P., Stallings, C.E., Abubakar, R.A., Mojahed, N., Guha, S., Abou-Jabal, D., and Ellsworth, B.S. (2025). The interplay between FOXO1 and glucocorticoid signaling in promoting the terminal differentiation of somatotropes. Mol. Cell. Endocrinol. 606, 112573. 10.1016/j.mce.2025.112573.

108. Klug, S.T., Ellestad, L.E., and Porter, T.E. (2025). Pituitary-Targeted Knockout of Glucocorticoid Receptors Disrupts Growth Hormone Expression During Embryonic Development. Endocrinology 166, bqaf119. 10.1210/endocr/bqaf119.

109. Miao, Y., Li, C., Guo, J., Wang, H., Gong, L., Xie, W., and Zhang, Y. (2019). Identification of a novel somatic mutation of POU6F2 by whole-genome sequencing in prolactinoma. Mol. Genet. Genomic Med. 7, e1022. 10.1002/mgg3.1022.

110. Schep, A.N., Wu, B., Buenrostro, J.D., and Greenleaf, W.J. (2017). chromVAR: inferring transcription-factor-associated accessibility from single-cell epigenomic data. Nat. Methods 14, 975–978. 10.1038/nmeth.4401.

111. Harris, H.K., Nakayama, T., Lai, J., Zhao, B., Argyrou, N., Gubbels, C.S., Soucy, A., Genetti, C.A., Suslovitch, V., Rodan, L.H., et al. (2021). Disruption of RFX family transcription factors causes autism, attention-deficit/hyperactivity disorder, intellectual disability, and dysregulated behavior. Genet. Med. 23, 1028–1040. 10.1038/s41436-021-01114-z.

112. Martinez-Mayer, J., Vishnopolska, S., Perticarari, C., Iglesias Garcia, L., Hackbartt, M., Martinez, M., Zaiat, J., Jacome-Alvarado, A., Braslavsky, D., Keselman, A., et al. (2024). Exome Sequencing Has a High Diagnostic Rate in Sporadic Congenital Hypopituitarism and Reveals Novel Candidate Genes. J. Clin. Endocrinol. Metab. 109, 3196–3210. 10.1210/clinem/dgae320.

113. Bashamboo, A., Bignon-Topalovic, J., Moussi, N., McElreavey, K., and Brauner, R. (2017). Mutations in the Human ROBO1 Gene in Pituitary Stalk Interruption Syndrome. J. Clin. Endocrinol. Metab. 102, 2401–2406. 10.1210/jc.2016-1095.

114. Dateki, S., Watanabe, S., Mishima, H., Shirakawa, T., Morikawa, M., Kinoshita, E., Yoshiura, K., and Moriuchi, H. (2019). A homozygous splice site ROBO1 mutation in a patient with a novel syndrome with combined pituitary hormone deficiency. J. Hum. Genet. 64, 341–346. 10.1038/s10038-019-0566-8.

115. Liu, Z., Chen, X., Liu, Z., and Chen, X. (2020). A Novel Missense Mutation in Human Receptor Roundabout-1 (ROBO1) Gene Associated with Pituitary Stalk Interruption Syndrome. J. Clin. Res. Pediatr. Endocrinol. 10.4274/jcrpe.galenos.2019.2018.0309.

116. Shamkiran, Sheshadri T., and Anil, M.U. (2025). Pituitary stalk interruption with multiple pituitary hormone deficiencies associated with a roundabout guidance receptor 1 gene mutation. J. Pediatr. Endocrinol. Diabetes 4, 139–142. 10.25259/JPED_41_2024.

117. Good, D.J., Porter, F.D., Mahon, K.A., Parlow, A.F., Westphal, H., and Kirsch, I.R. (1997). Hypogonadism and obesity in mice with a targeted deletion of the Nhlh2 gene. Nat. Genet. 15, 397–401. 10.1038/ng0497-397.

118. Leon, S., Talbi, R., McCarthy, E.A., Ferrari, K., Fergani, C., Naule, L., Choi, J.H., Carroll, R.S., Kaiser, U.B., Aylwin, C.F., et al. (2021). Sex-specific pubertal and metabolic regulation of Kiss1 neurons via Nhlh2. eLife 10, e69765. 10.7554/eLife.69765.

119. Cogliati, T., Delgado-Romero, P., Norwitz, E.R., Guduric-Fuchs, J., Kaiser, U.B., Wray, S., and Kirsch, I.R. (2007). Pubertal Impairment in Nhlh2 Null Mice Is Associated with Hypothalamic and Pituitary Deficiencies. Mol. Endocrinol. 21, 3013–3027. 10.1210/me.2005-0337.

120. Stinnett, G.S., Westphal, N.J., and Seasholtz, A.F. (2015). Pituitary CRH-binding protein and stress in female mice. Physiol. Behav. 150, 16–23. 10.1016/j.physbeh.2015.02.050.

121. Gammie, S.C., Seasholtz, A.F., and Stevenson, S.A. (2008). Deletion of corticotropin-releasing factor binding protein selectively impairs maternal, but not intermale aggression. Neuroscience 157, 502–512. 10.1016/j.neuroscience.2008.09.026.

122. Jin, K., Yao, Z., van Velthoven, C.T.J., Kaplan, E.S., Glattfelder, K., Barlow, S.T., Boyer, G., Carey, D., Casper, T., Chakka, A.B., et al. (2025). Brain-wide cell-type-specific transcriptomic signatures of healthy ageing in mice. Nature 638, 182–196. 10.1038/s41586-024-08350-8.

123. Wilkinson, M.D., Dumontier, M., Aalbersberg, Ij.J., Appleton, G., Axton, M., Baak, A., Blomberg, N., Boiten, J.-W., da Silva Santos, L.B., Bourne, P.E., et al. (2016). The FAIR Guiding Principles for scientific data management and stewardship. Sci. Data 3, 160018. 10.1038/sdata.2016.18.

124. Gálvez-Merchán, Á., Min, K.H. (Joseph), Pachter, L., and Booeshaghi, A.S. (2023). Metadata retrieval from sequence databases with ffq. Bioinformatics 39, btac667. 10.1093/bioinformatics/btac667.

125. Sullivan, D.K., Min, K.H. (Joseph), Hjörleifsson, K.E., Luebbert, L., Holley, G., Moses, L., Gustafsson, J., Bray, N.L., Pimentel, H., Booeshaghi, A.S., et al. (2024). kallisto, bustools, and kb-python for quantifying bulk, single-cell, and single-nucleus RNA-seq. bioRxiv, 2023.11.21.568164. 10.1101/2023.11.21.568164.

126. Bray, N.L., Pimentel, H., Melsted, P., and Pachter, L. (2016). Near-optimal probabilistic RNA-seq quantification. Nat. Biotechnol. 34, 525–527. 10.1038/nbt.3519.

127. Mathys, H., Davila-Velderrain, J., Peng, Z., Gao, F., Mohammadi, S., Young, J.Z., Menon, M., He, L., Abdurrob, F., Jiang, X., et al. (2019). Single-cell transcriptomic analysis of Alzheimer’s disease. Nature 570, 332–337. 10.1038/s41586-019-1195-2.

128. Mendelev, N., Zamojski, M., Amper, M.A.S., Cheng, W.S., Pincas, H., Nair, V.D., Zaslavsky, E., Sealfon, S.C., and Ruf-Zamojski, F. (2022). Multi-omics profiling of single nuclei from frozen archived postmortem human pituitary tissue. STAR Protoc. 3, 101446. 10.1016/j.xpro.2022.101446.

129. Zhang, Z., Zamojski, M., Smith, G.R., Willis, T.L., Yianni, V., Mendelev, N., Pincas, H., Seenarine, N., Amper, M.A.S., Vasoya, M., et al. (2022). Single nucleus transcriptome and chromatin accessibility of postmortem human pituitaries reveal diverse stem cell regulatory mechanisms. Cell Rep. 38, 110467. 10.1016/j.celrep.2022.110467.

130. Wolf, F.A., Angerer, P., and Theis, F.J. (2018). SCANPY: large-scale single-cell gene expression data analysis. Genome Biol. 19, 15. 10.1186/s13059-017-1382-0.

131. Sheng, C., Lopes, R., Li, G., Schuierer, S., Waldt, A., Cuttat, R., Dimitrieva, S., Kauffmann, A., Durand, E., Galli, G.G., et al. (2022). Probabilistic machine learning ensures accurate ambient denoising in droplet-based single-cell omics. Preprint at bioRxiv, 10.1101/2022.01.14.476312 10.1101/2022.01.14.476312.

132. Zhang, A.W., O’Flanagan, C., Chavez, E.A., Lim, J.L.P., Ceglia, N., McPherson, A., Wiens, M., Walters, P., Chan, T., Hewitson, B., et al. (2019). Probabilistic cell-type assignment of single-cell RNA-seq for tumor microenvironment profiling. Nat. Methods 16, 1007–1015. 10.1038/s41592-019-0529-1.

133. Zhang, K., Zemke, N.R., Armand, E.J., and Ren, B. (2024). A fast, scalable and versatile tool for analysis of single-cell omics data. Nat. Methods 21, 217–227. 10.1038/s41592-023-02139-9.

134. Zhang, Y., Liu, T., Meyer, C.A., Eeckhoute, J., Johnson, D.S., Bernstein, B.E., Nusbaum, C., Myers, R.M., Brown, M., Li, W., et al. (2008). Model-based Analysis of ChIP-Seq (MACS). Genome Biol. 9, R137. 10.1186/gb-2008-9-9-r137.

135. Amemiya, H.M., Kundaje, A., and Boyle, A.P. (2019). The ENCODE Blacklist: Identification of Problematic Regions of the Genome. Sci. Rep. 9, 9354. 10.1038/s41598-019-45839-z.

136. Martens, L.D., Fischer, D.S., Yépez, V.A., Theis, F.J., and Gagneur, J. (2024). Modeling fragment counts improves single-cell ATAC-seq analysis. Nat. Methods 21, 28–31. 10.1038/s41592-023-02112-6.

137. Gayoso, A., Lopez, R., Xing, G., Boyeau, P., Valiollah Pour Amiri, V., Hong, J., Wu, K., Jayasuriya, M., Mehlman, E., Langevin, M., et al. (2022). A Python library for probabilistic analysis of single-cell omics data. Nat. Biotechnol. 40, 163–166. 10.1038/s41587-021-01206-w.

138. Clarke, Z.A., and Bader, G.D. (2024). MALAT1 expression indicates cell quality in single-cell RNA sequencing data. Preprint at bioRxiv, 10.1101/2024.07.14.603469 10.1101/2024.07.14.603469.

139. Badia-I-Mompel, P., Vélez Santiago, J., Braunger, J., Geiss, C., Dimitrov, D., Müller-Dott, S., Taus, P., Dugourd, A., Holland, C.H., Ramirez Flores, R.O., et al. (2022). decoupleR: ensemble of computational methods to infer biological activities from omics data. Bioinforma. Adv. 2, vbac016. 10.1093/bioadv/vbac016.

140. Law, C.W., Chen, Y., Shi, W., and Smyth, G.K. (2014). voom: precision weights unlock linear model analysis tools for RNA-seq read counts. Genome Biol. 15, R29. 10.1186/gb-2014-15-2-r29.

141. Hoffmann, E.S., Hombrecher, L., Diks, I.F., Flotho, M., Keller, A., and Grandke, F. (2025). PBMCpedia: A Harmonized PBMC scRNA-seq Database With Unified Mapping and Enhanced Celltype Annotation. Preprint at bioRxiv, 10.1101/2025.08.06.668843 10.1101/2025.08.06.668843.

142. Pedregosa, F., Varoquaux, G., Gramfort, A., Michel, V., Thirion, B., Grisel, O., Blondel, M., Prettenhofer, P., Weiss, R., Dubourg, V., et al. (2011). Scikit-learn: Machine Learning in Python. J. Mach. Learn. Res. 12, 2825–2830.

143. Schindelin, J., Arganda-Carreras, I., Frise, E., Kaynig, V., Longair, M., Pietzsch, T., Preibisch, S., Rueden, C., Saalfeld, S., Schmid, B., et al. (2012). Fiji: an open-source platform for biological-image analysis. Nat. Methods 9, 676–682. 10.1038/nmeth.2019.

