## Supplemental Figures 1-6 for "Consensus Pituitary Atlas, a scalable resource for annotation, novel marker discovery and analyses in pituitary gland research"

### Supplemental information - Kövér *et al.*

#### Document S1. Figures S1–S7

**Supplementary Table 1: Curation of single-cell datasets**
**Supplementary Table 2: Consensus Chromatin Landscape peaks**
**Supplementary Table 3: Table of all cell typing markers**
**Supplementary Table 4: Table of all sex-biased genes**
**Supplementary Table 5: Table of all age-dependent genes**
**Supplementary Table 6: Table of all lineage marker genes**
**Supplementary Table 7: Table of all low expression markers**
**Supplementary Table 8: Table of enrichment results for stem cells**
**Supplementary Table 9: Table of all signaling results**
**Supplementary Table 10: Table of all lineage marker peaks**
**Supplementary Table 11: Table of all TF hits**
**Supplementary Table 12: Curation of bulk datasets**

#### Supplementary Figure Legends

**Supplementary Figure 1**

(A) Double bar plot of the number of captured fragments using per-dataset called peaks (gray - lower) versus using the Consensus Chromatin Landscape (blue - higher).

(B) (Left) Box plot of cell numbers per cell type across single-nucleus ATAC-seq datasets (each dot is a pseudobulk sample), showing the overrepresentation of somatotrophs and lactotrophs. Bar heights are showing the median values. (Middle) Box plot of the number of peaks with ≥1 fragment captured per cell type across single-nucleus ATAC-seq datasets (each dot is a pseudobulk sample). This shows that despite the overrepresentation of somatotrophs and lactotrophs, the peaks represent all cell types in a balanced way. Bar heights are showing the median values. (Right) Box plot of the average fragment count per cell type across single-nucleus ATAC-seq datasets (each dot is a pseudobulk sample). This shows that despite the overrepresentation of somatotrophs and lactotrophs, the peaks represent all cell types in balanced way. Bar heights are showing the median values.

(C) (Left) Stacked bar plot of cell type composition across all ATAC samples and (Right) mean cell type composition of male and female samples.

**Supplementary Figure 2**

(A) Violin plots of log1p total counts and log1p number of genes in doublets and real cells (as identified by the Doublet Model) in the newly generated P4 male multiome dataset.

(B) Panel of UMAPs for *Tbx19*-KO sample.[34] Cell types were assigned by the Cell Type Model. The various genes shown are melanotroph markers determined using the CPA.

(C) Row-normalized (giving recall percentages on the diagonal) confusion matrix of Cell Type Model predicted labels (X axis) and true cell type labels (Y axis). These results were derived from the chromatin accessibility Cell Type Model.

**Supplementary Figure 3**

(A) Volcano plot of sex-biased genes from reanalysis of male/female gonadotroph bulk RNA-seq samples,[64] colored according to sex-bias in CPA. Gonadotroph sex-biased genes *Fshb*, *Gpr101* and *Grem1* are highlighted in red.

(B) Volcano plot of sex-biased genes from reanalysis of male/female corticotroph bulk RNA-seq samples,[65] colored according to sex-bias in CPA.

(C) Box plot of *Gpr101* expression across cell types and sexes (blue: male; orange: female).

(D) Box plot of *Gata2, Grem1, and Fshb* expression in gonadotrophs showing sex-biased expression (blue: male; orange: female).

(E) Horizontal strip plot of $\log_2$ fold-changes in Ctrl vs AX and Ctrl vs AX/GX comparisons (blue: upregulated; orange: downregulated).

**Supplementary Figure 4**

(A) Scatter plot of immune cell proportions across all RNA samples versus $\log_{10}$ age. The data suggest an increase in immune cells with age.

(B) Scatter plots of gene expression versus age (in $\log_{10}$ days) of 10 selected statistically significant changing signaling genes in stem cells.

(C) Scatter plots of gene expression versus age (in $\log_{10}$ days) of statistically significant changing TF genes in stem cells.

(D) Immunofluorescence staining against LEF1 (red) and SOX2 (green) in anterior pituitary parenchyma of mice at P3, P15 and P56. Scale bars 50 µm.

(E) Quantification of LEF1+SOX2+ double expressing cells over SOX2+ cells in the parenchyma (P, green) and marginal zone (MZ, blue) in P3, P15 and P56 animals. Each dot is a biological replicate. Two-sided paired T-test, *p*-values indicated.

**Supplementary Figure 5**

(A) Dot plot of 20 genes (chosen at random from 576) identified as stem cell markers in all three individual datasets examined in Figure 5B, but not using the CPA. Dot plot shows expression is not selective for stem cells.

(B) Line plot showing the same analysis as presented in Figure 5B, except repeated for all lineage comparisons, indicating the number of genes that fall within the various parts of the Venn diagram. The highest discrepancy is in the Corticotroph/Melanotroph comparison, where 53 genes are detected as differentially expressed in the CPA, compared to 2501 genes using individual datasets.

(C) Dot plot of all Eph-Ephrin signaling genes.

(D) Dot plot of stem cell-specific Laminin signaling genes.

(E) Summary of LIANA+ results. Blue shades represent various degrees of specificity ($-\log_{10}$ specificity_rank), such that the darker colors represent more specific interactions.

(F) Dot plot of BMP6 interaction genes, showing the possible signaling interaction between stem cells and other cells of the pituitary.

(G) Dot plot of FGF1-FGFR1/2 interaction genes, showing stem cells as the main *Fgf* source, in addition to pituicytes and corticotrophs. The main receptors *Fgfr1* and *Fgfr2* are highly expressed across cells, though *Fgfr1* is primarily specific to stem cells from anterior pituitary cell types.

(H) Dot plot of TGFB2-TGFBR interaction genes, showing stem cells as the main source of signals.

**Supplementary Figure 6**

(A) Schematic of the number of RNA-only hit TFs in the differentiation hierarchy of the pituitary lineage. Purple numbers mark upregulated TFs, while orange numbers mark downregulated TFs at each branch point.

(B) Table of all RNA-only hits. Upregulated in purple, downregulated in orange.

(C) Box plot of *Nhlh2* expression, revealing female bias.

(D) Bar plot of TF motif fold-enrichment for each Multimodal hit TF in stem cells.

(E) Heatmap of the number of peaks marked by the motif of a stem cell Multimodal TF falling into various annotated regions. This plot offers an explanation as to why KLF motifs appear to be separated from the binding sites of other TFs. Specifically, KLF motifs appear enriched in proximal regulatory regions compared to other TFs.

(F) Strip plot of motif to TSS distances, with a threshold at 5kb. The X-axis is ordered according to the percentage of motif distances below 5kb. This further supports the hypothesis that KLF motifs are enriched in promoter proximal regions compared to other TF motifs.

Supplementary Figure 1 - Kövér et al.

A

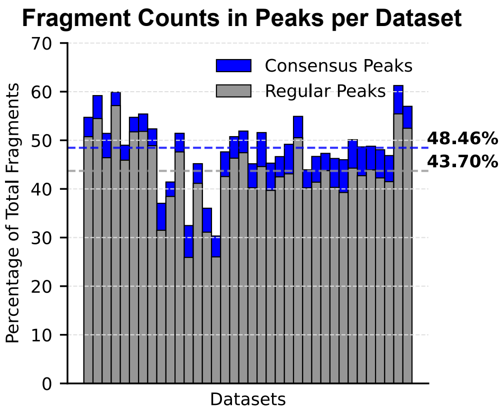


B

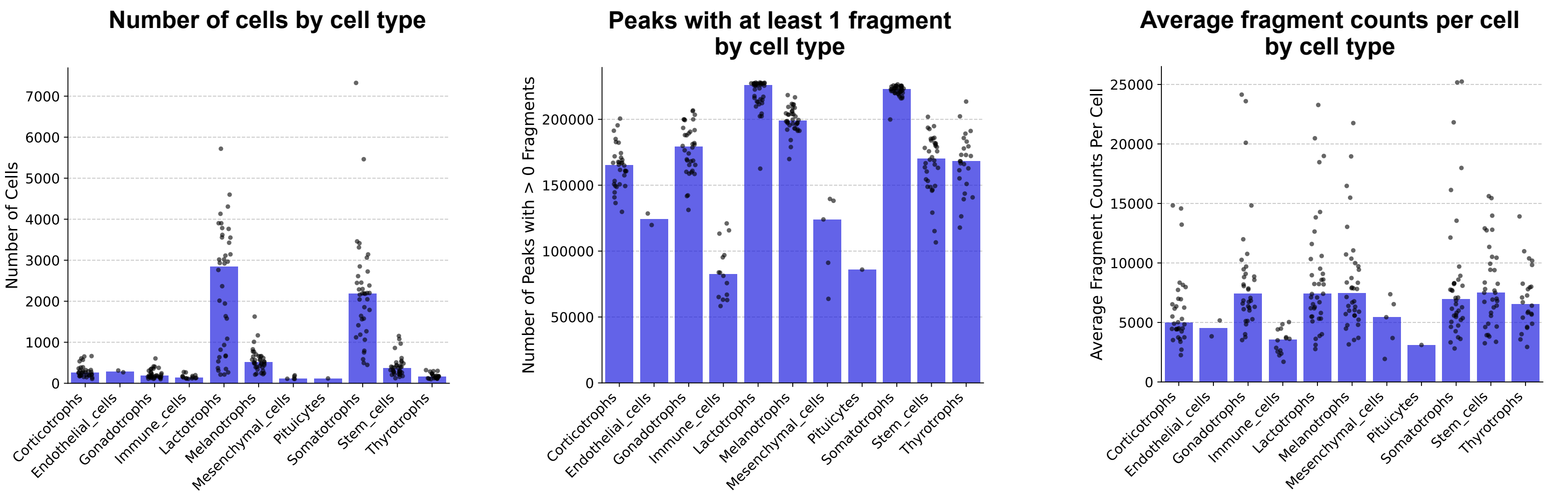


C

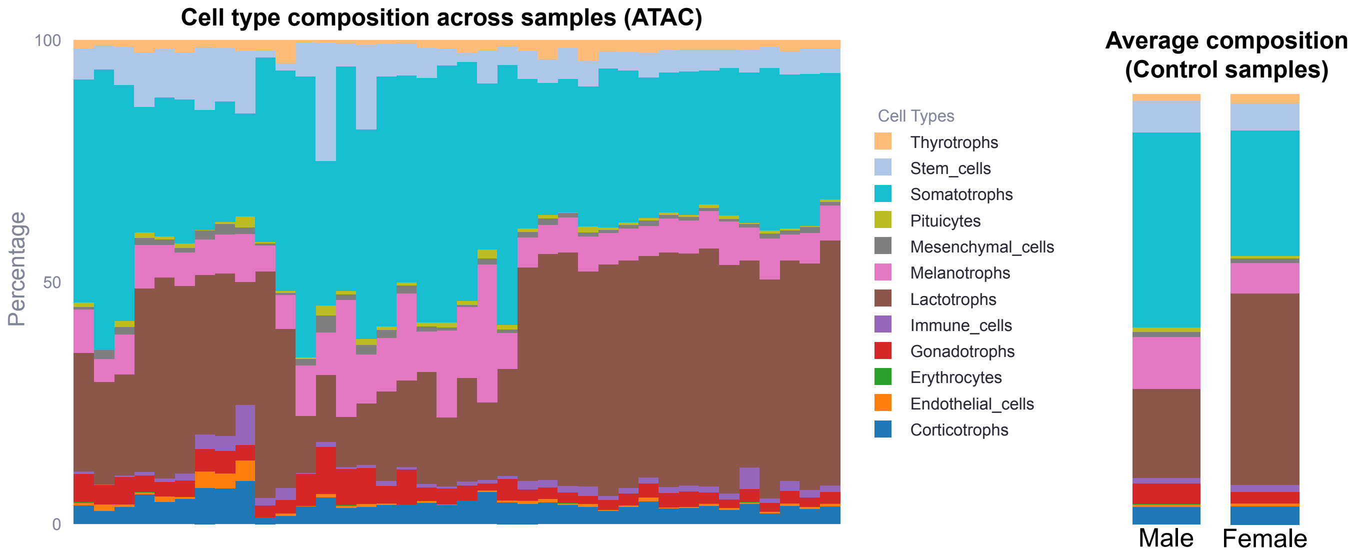

Supplementary Figure 2 - Kövér et al.

**A**

**Predicted doublets exhibit higher counts and number of genes**

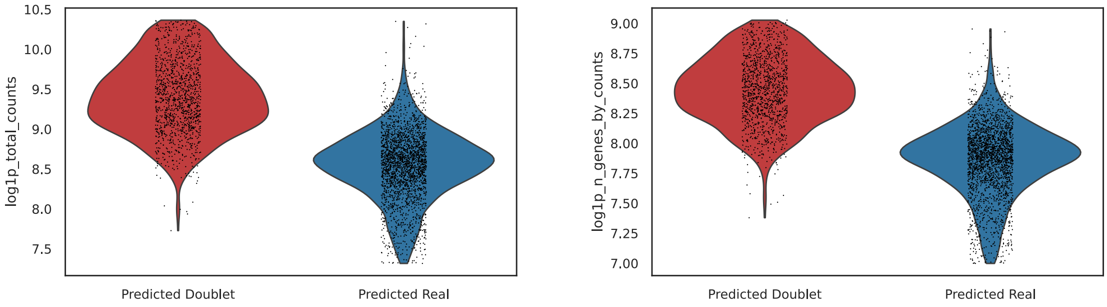


**B**

**Melanotroph markers highlight the predicted melanotroph population in the *Tbx19*-KO dataset**

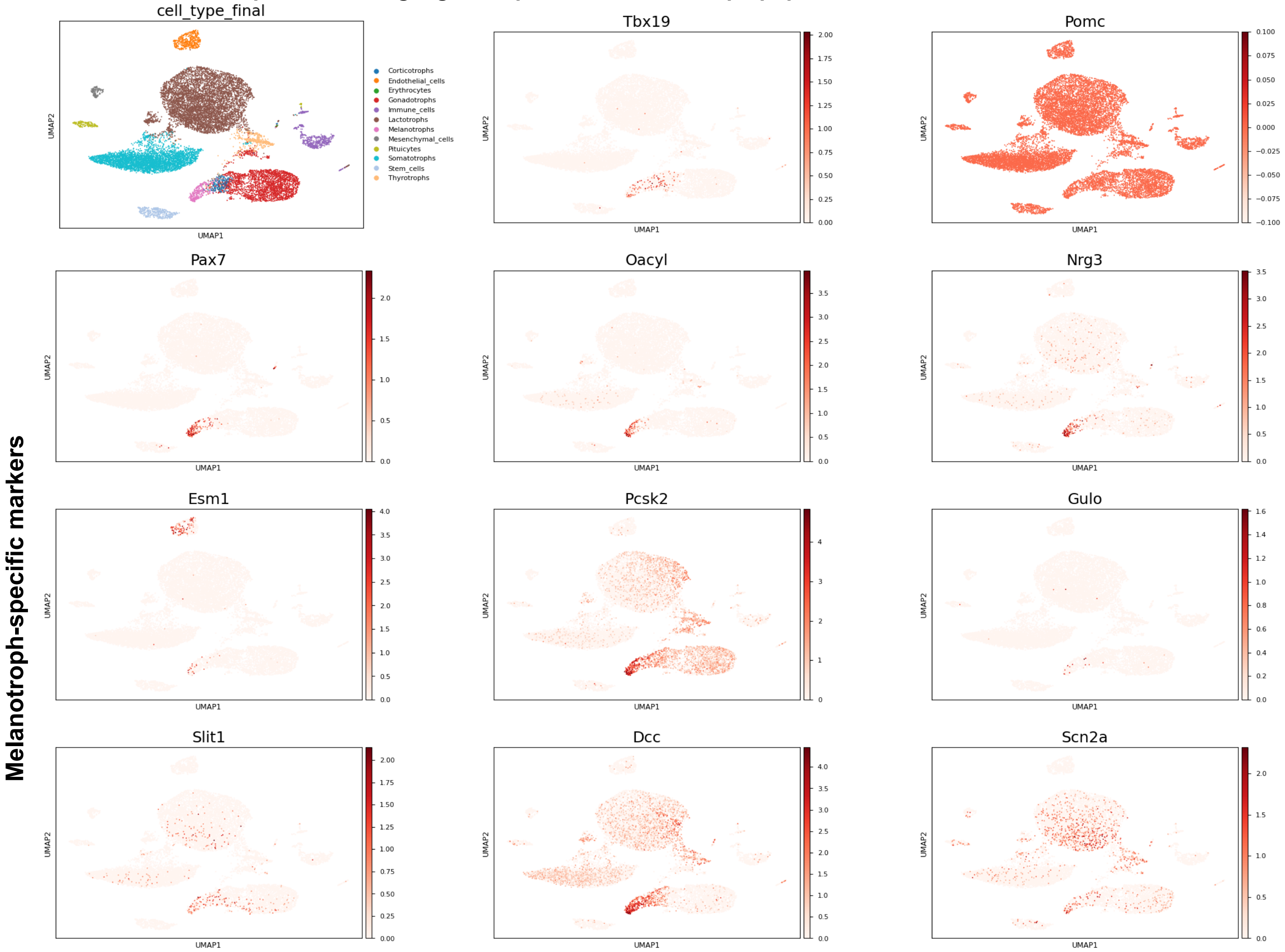


**C**

**Row-normalised ("Recall") Confusion Matrix - ATAC**

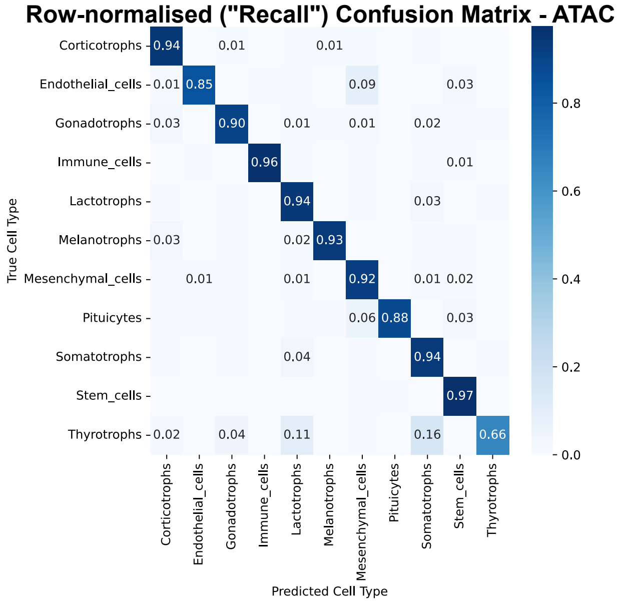

Supplementary Figure 3 - Kövér et al.

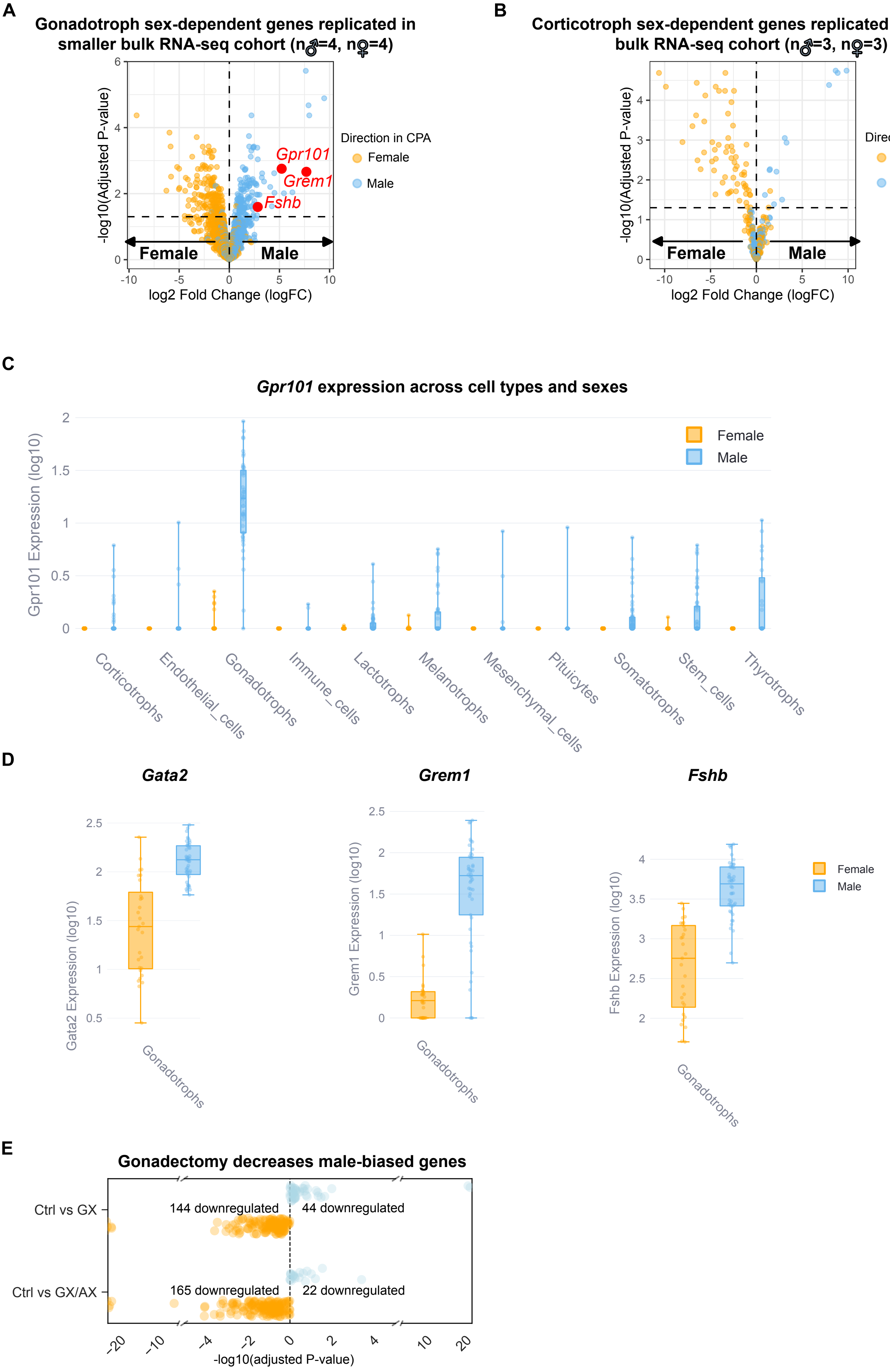

Supplementary Figure 4 - Kövér et al.

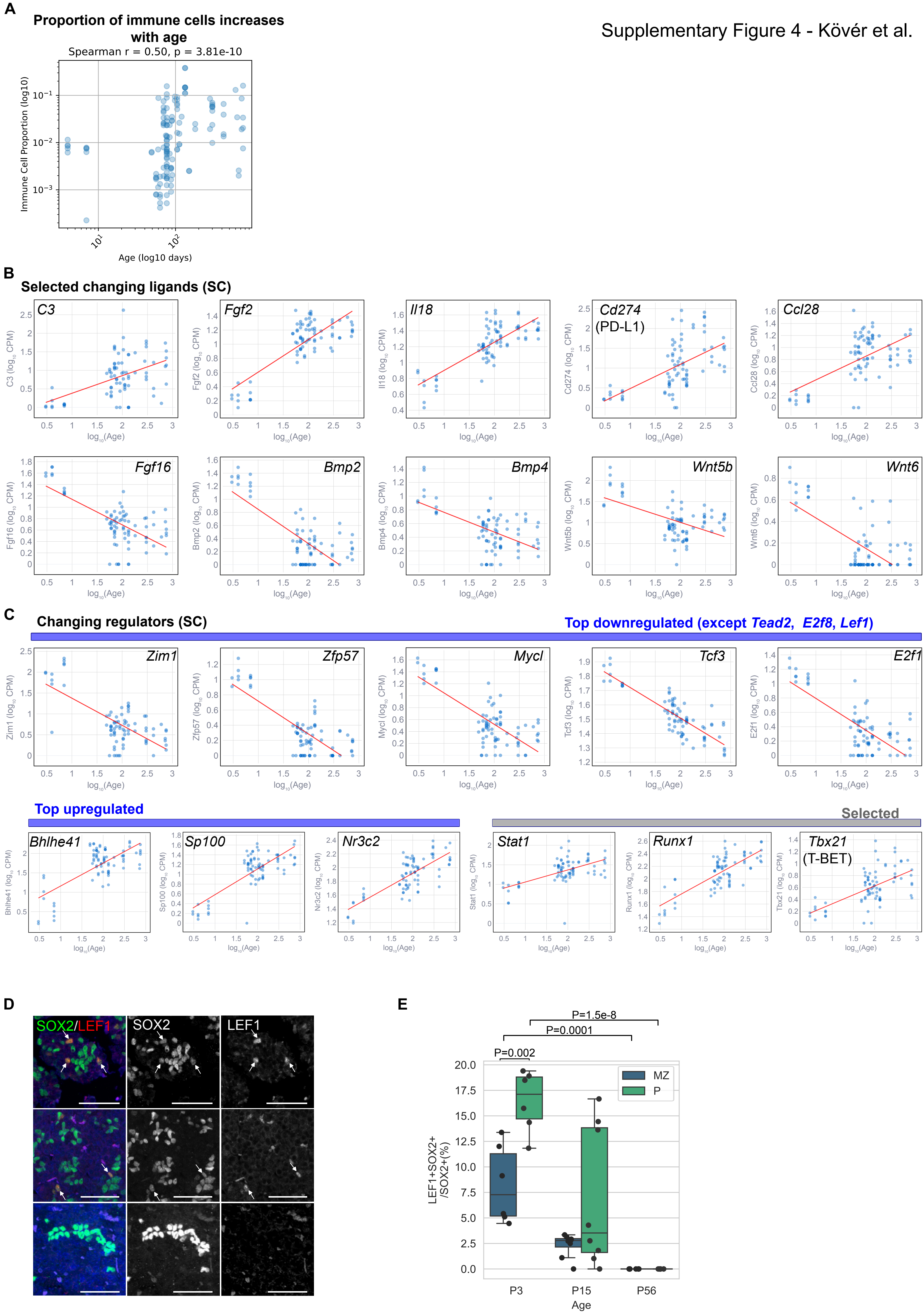

Supplementary Figure 5
Kövér et al.

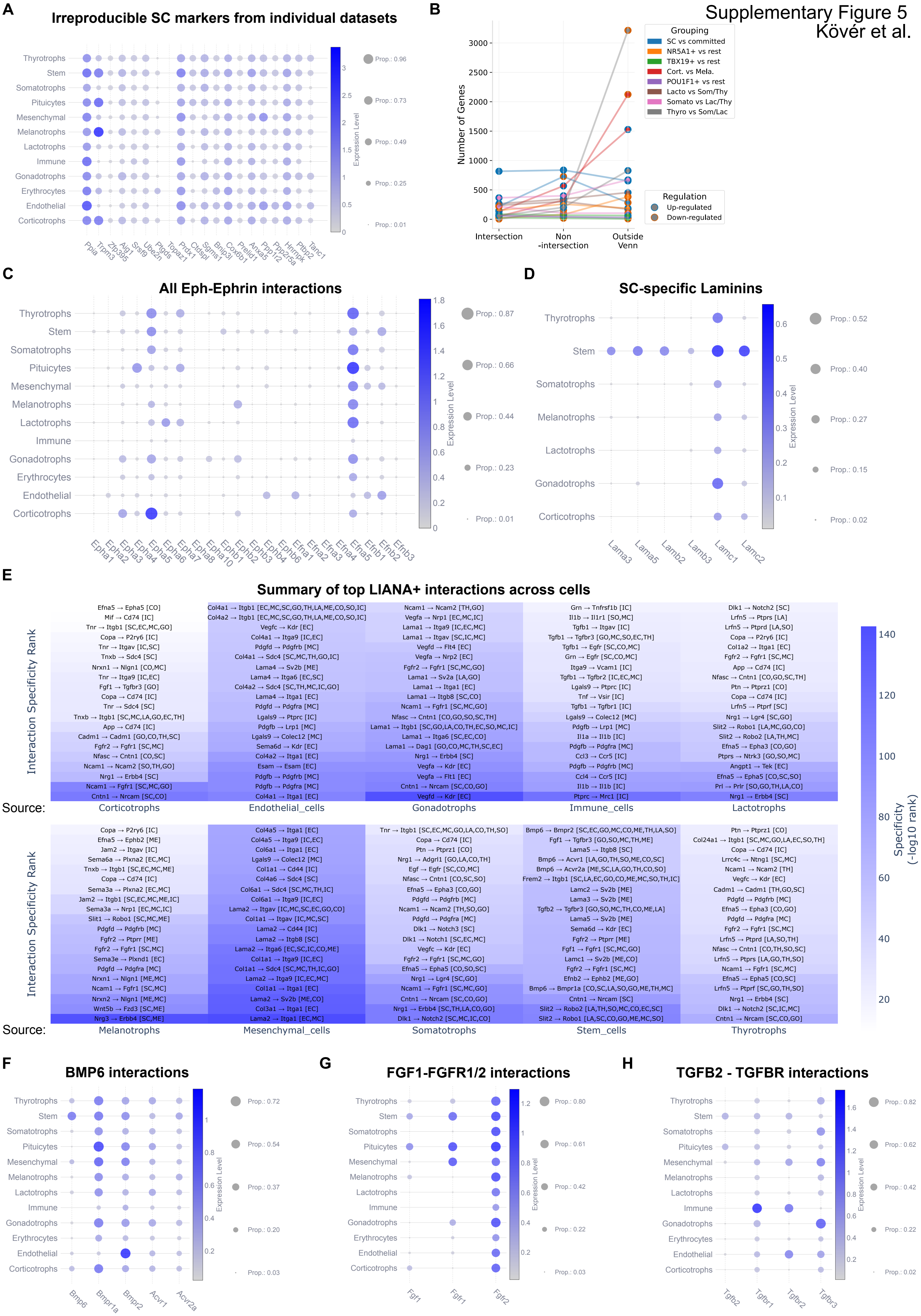

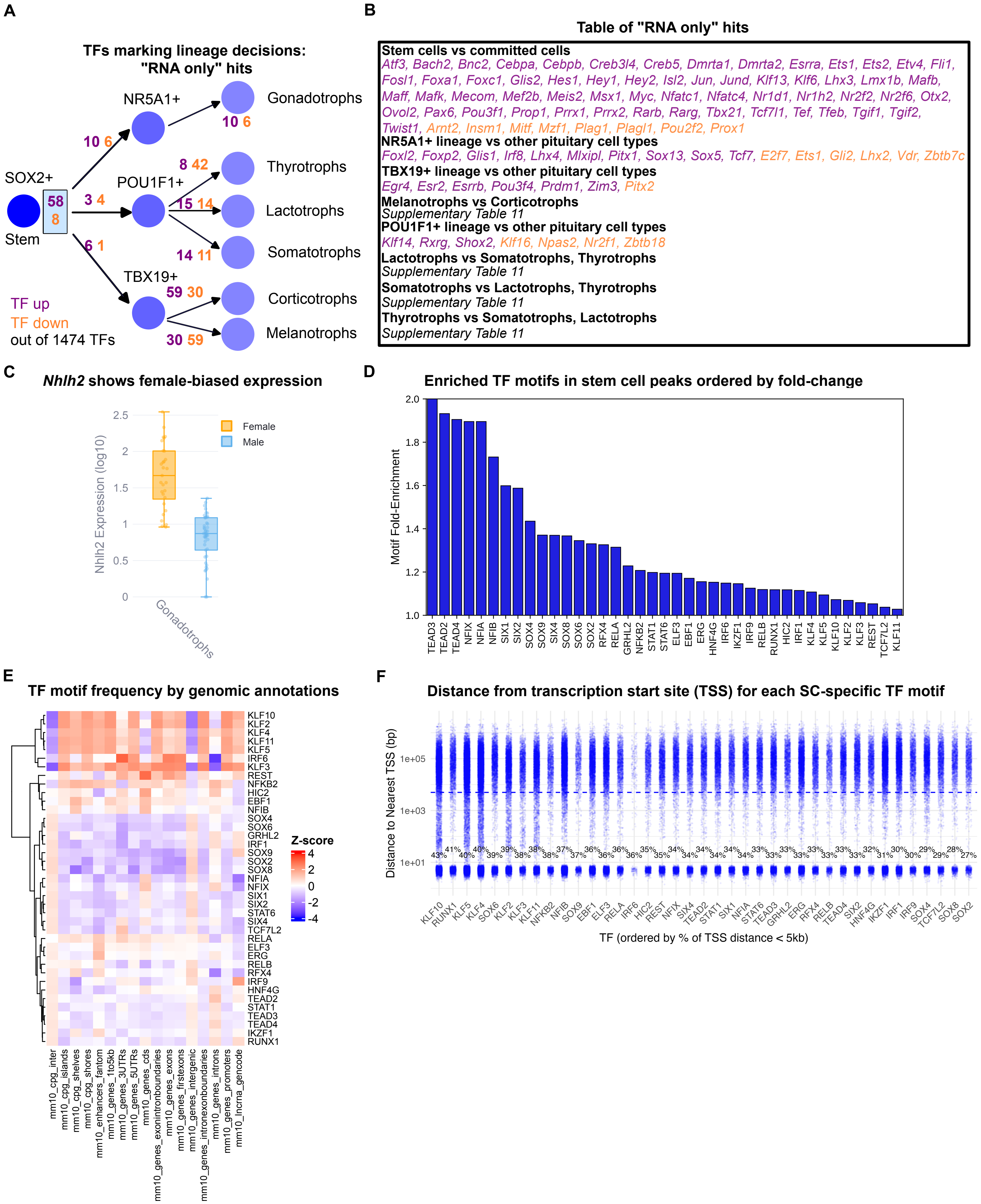


Supplementary Figure 6 - Kövér et al.
